# The representational space of observed actions

**DOI:** 10.1101/592071

**Authors:** Raffaele Tucciarelli, Moritz F. Wurm, Elisa Baccolo, Angelika Lingnau

## Abstract

Categorizing and understanding other people’s actions is a key human capability. Whereas there exists a growing literature regarding the organization of objects, the representational space underlying the organization of observed actions remain largely unexplored. Here we examined the organizing principles of a large set of actions and the corresponding neural representations. Using multiple-regression representational similarity analysis of fMRI data, in which we accounted for variability due to major action-related features (body parts, scenes, movements, objects), we found that the semantic dissimilarity structure was best captured by patterns of activation in the lateral occipitotemporal cortex (LOTC) and the left posterior inferior parietal lobe (IPL). Together, our results demonstrate that the organization of observed actions in the LOTC and the IPL resembles the organizing principles used by participants to classify actions behaviorally, in line with the view that these regions are crucial for accessing the meaning of actions.

---

Humans can perform a striking number of different types of actions, from hammering a nail to performing open heart surgery. However, most of what we know about the way we control and recognize actions is based on a rich literature on prehension movements in humans and non-human primates. This literature revealed a widespread network of fronto-parietal regions, with a preference along the dorso-medial stream and the dorso-lateral stream for reaching and grasping movements, respectively (for reviews see e.g.^1–3^). Less is known regarding the organization of more complex actions (for exceptions, see^4–6^). According to which principles are different types of actions organized in the brain, and do these principles help us understand how we are able to tell that two actions, e.g. running and riding a bike, are more similar to each other than two other actions, e.g. driving a bike and reading a book? Are observed actions that we encounter on a regular basis organized according to higher-level semantic categorical distinctions (e.g. between locomotion, object manipulation, and communication actions and further overarching organizational dimensions)? Note that higher-level semantic categories often covary with more basic action-related aspects of perceived action scenes (e.g. locomotion actions often take place outdoors and do not involve objects, communicative actions often involve mouth/lip movements, etc.). Disentangling these levels neurally presents an analytical challenge that has not been addressed so far.

A number of recent studies used multivariate pattern analysis (MVPA)^7^ to examine which brain areas are capable to distinguish between different observed actions (e.g. opening vs closing, slapping vs lifting an object, or cutting vs peeling)^6,8–12^. The general results that emerged from these studies is that it is possible to distinguish between different actions on the basis of patterns of brain activation in the lateral occipito-temporal cortex (LOTC), the inferior parietal lobe (IPL) and the ventral premotor cortex (PMv). In line with this view, LOTC has been shown to contain action-related object properties^13^. LOTC and IPL, but not the PMv, furthermore showed a generalization across the way in which these actions were performed (e.g. performing the same action with different kinematics), suggesting that these areas represent actions at more general levels and thus possibly the meaning of the actions. However, previous studies could not unambiguously determine what kind of information was captured from observed actions: movement trajectories and body postures^6,11,12^, certain action precursors at high levels of generality^9^ (e.g. object state change), or more complex semantic aspects that go beyond the basic constituents of perceived actions and that represent the meaning of actions at higher integratory levels. In the latter case, the LOTC and the IPL should also reflect the semantic similarity structure of a wide range of actions: Running shares more semantic aspects with riding a bike than with reading; therefore, semantic representations of running and riding a bike should be more similar with each other than with the semantic representation of reading.

To test the prediction that the LOTC and the IPL reflect the semantic similarity structure of a wide range of actions, we carried out a behavioral and an fMRI study in the same group of participants. In the behavioral experiment, we used inverse multidimensional scaling (MDS)^14^ to examine the structure of the similarity in meaning (which we will refer to as *semantic similarity* in the remainder of this paper) of a range of different actions (Figure 1). Moreover, to control for more basic constituents of actions that often covary with action semantics, we carried out inverse MDS for *action-related* aspects that typically constitute the basic perceptual cues in naturalistic actions, namely, body parts, scenes, movement kinematics, and objects. In the fMRI study, we examined which brain regions capture the semantic similarity structure determined in the behavioral experiment, using representational similarity analysis^15^ (RSA). Moreover, to examine which brain areas capture semantic similarity over and beyond the other action-related aspects we examined, we carried out a multiple-regression RSA for each of the behavioral models while accounting for the remaining models.

**Figure 1.**
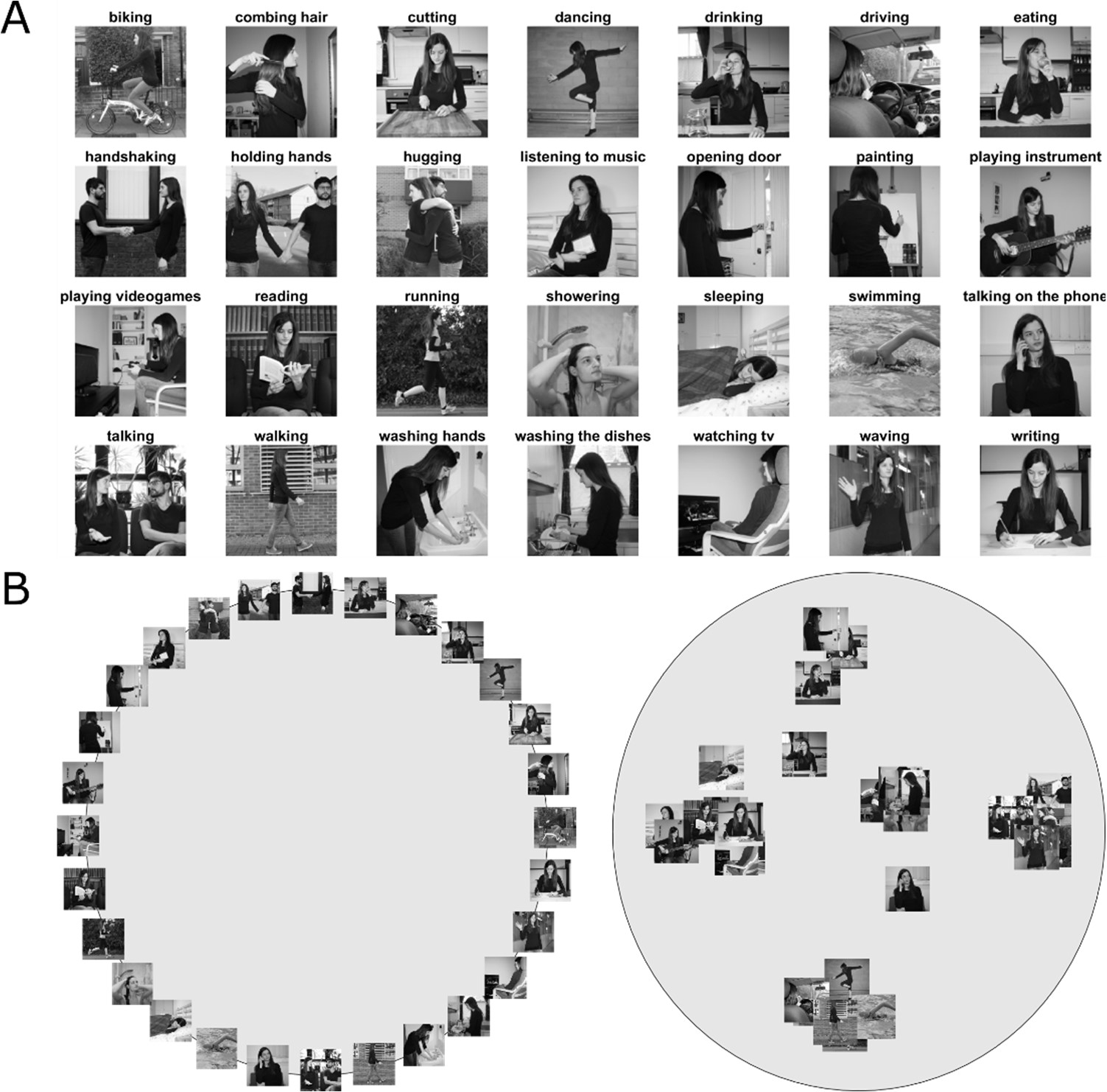
Stimuli and behavioral task. A) Stimuli used in the behavioral and the fMRI experiment, depicting static images of 28 everyday actions. To increase perceptual variability, actions were shown from different viewpoints, in different scenes, using two different actors (see text for details) and different objects (for actions involving an object). For a full set of stimuli used in the fMRI experiment, see Supplementary Information, Figure S1. B) Illustration of the behavioral experiment used for inverse multidimensional scaling. In the first trial of the experiment, participants were presented with an array of images arranged on the circumference of a grey circle (left panel). In each subsequent trial, an adaptive algorithm determined a subset of actions that provided optimal evidence for the pairwise dissimilarity estimates (see ^14^ and Methods, for details). In different parts of the experiment, participants were asked to rearrange the images according to their perceived similarity with respect to a specific aspect of the action, namely, their meaning (or semantics), the body part(s) involved, the scene/ context in which the action typically takes place, movement kinematics, and objects involved in the action. Right panel: Example arrangement resulting from the semantic task of one representative participant. Using inverse multidimensional scaling, we derived a behavioral model (see Figure 2) from this arrangement, individually for each participant, that we then used for the representational similarity analysis to individuate those brain regions that showed a similar representational geometry (for details, see Methods section).

## Results

### Behavioral

#### Inverse Multidimensional Scaling Experiment (Behavioral)

To investigate how similar participants judged the actions to be with regard to their meaning (referred to as *semantics* in the remainder of this paper) and action-related aspects (body parts, movement kinematics, objects, scenes), we extracted the pairwise Euclidean distances from the participants’ inverse MDS arrangements to obtain dissimilarity matrices (DSMs) for each participant and each behavioral model. Figure 2 shows DSMs for each model averaged across participants. For each model, we found significant (all p-values were smaller than p < 0.0001 and survived false discovery rate correction) inter-observer correlations, i.e., the individual DSMs significantly correlated with the average DSM of the remaining participants (mean leave-one-subject-out correlation coefficients [min – max individual correlation coefficients]; body: 0.57 [0.31 – 0.70]; scene: 0.63 [0.40 – 0.78]; movement: 0.47 [0.26 – 0.67]; object: 0.51 [0.22 – 0.71]; semantic: 0.61 [0.46 – 0.78]), suggesting that the participants’ arrangements were reliable and based on comparable principles.

**Figure 2.**
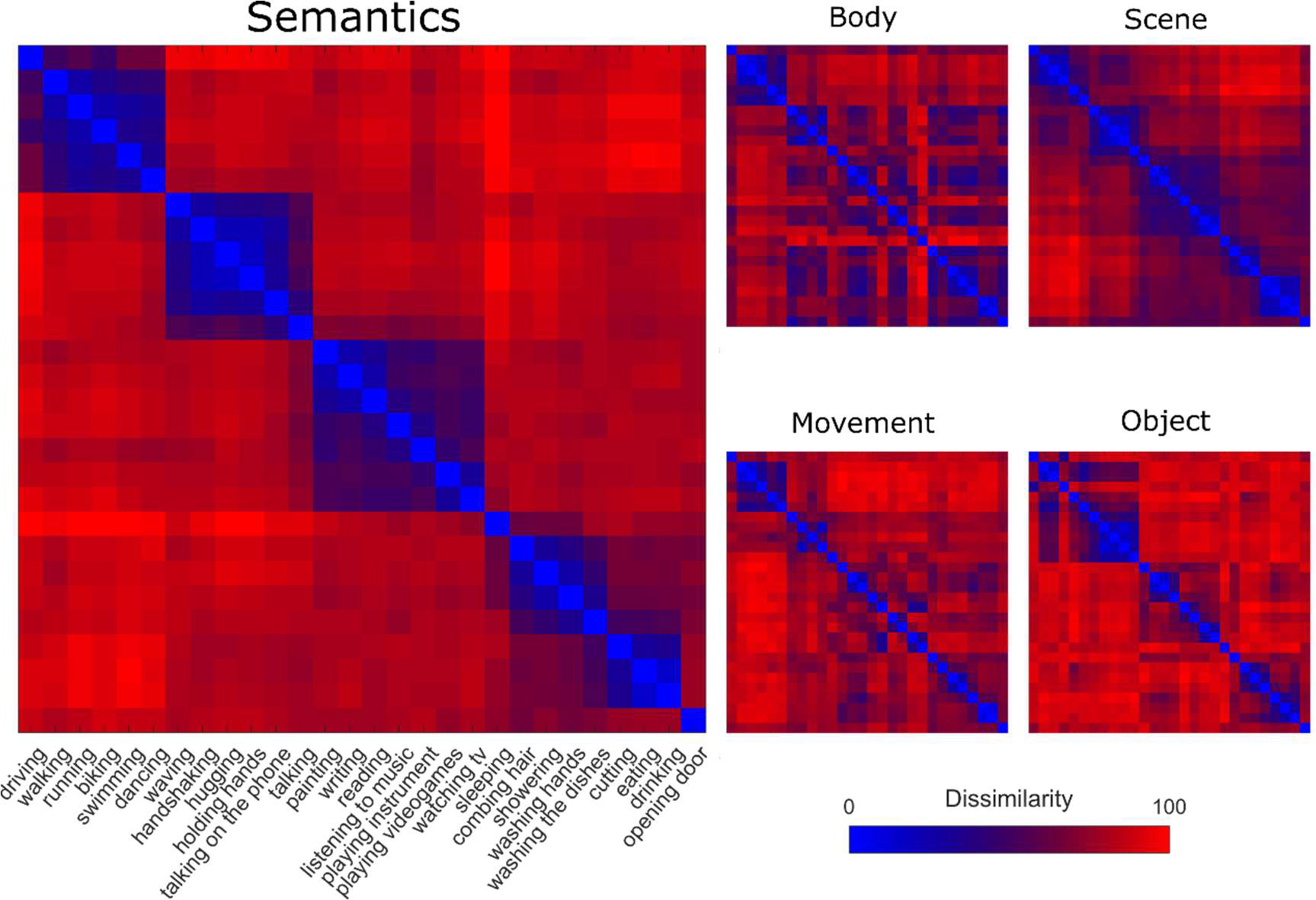
Behavioral Dissimilarity Matrices. Group dissimilarity matrices derived from inverse MDS using five different tasks (semantics, body, scene, movement, object). Bluish colors indicate high similarity between pairwise combinations of actions, whereas reddish colors indicate high dissimilarity. For ease of comparison, we used the ordering of the actions resulting from the semantic task for the remaining tasks (body, scene, movement, and object).

To examine correlations between models, we computed correlations between each pairwise comparison of DSMs, both averaged across participants (Supplementary Information, Figure S2A) and separately for each participant (Supplementary Information, Figure S2B, C). As can be seen, not surprisingly, there are modest correlations between the different models (in particular when averaged across participants). However, as shown in the Methods section on Multiple Regression RSA, the Variance Inflation Factor, computed for each participant, indicated a low risk of collinearity and thus justified the use of these models for multiple regression analysis.

To test for putative correlations of the semantic model with low-level image characteristics of the stimuli, we created a pixelwise similarity model by vectorizing each of the 28*12 (action*exemplar) = 336 action images, correlating each image with each other (resulting in a 336*336 correlation matrix), averaging the correlation coefficients across the 12 exemplars (resulting in a 28*28 matrix), and converting the correlation matrix into a (1 – r). There was no significant correlation between the pixelwise and semantic models (r= 0.038, p = 0.47). With regard to the action-related feature models, the body model and the movement model showed weak but significant correlations with the pixelwise model (r/p; object: −0.01/0.9, scene: 0.09/0.07, body: 0.11/0.04, movement: 0.1/0.05).

#### Principal Component Analysis (PCA) and Clustering analysis: K-means

To better characterize the dimensions along which the actions were organized and the clusters resulting from inverse MDS, we carried out a K-means clustering analysis and a Principal Component Analysis (PCA; see **Methods** for details). The Silhouette method (see **Methods** and Figure S3) revealed that the optimal number of clusters for the semantic task was six. As can be seen in Figure S4, the first three components account for the largest amount of variance. For ease of visualization, we show the first two components in Figure 3. A visualization of the first three principal components is shown in Figure S5. The analysis revealed clusters related to locomotion (e.g. biking, running), social/communicative actions (e.g. handshaking, talking on the phone), leisure-related actions (e.g. painting, reading), food-related actions (e.g. eating, drinking), and cleaning-related actions (e.g. showering, washing the dishes). Clusters obtained from the remaining models can be found in the Supplementary Information (Figure S6A-D).

**Figure 3.**
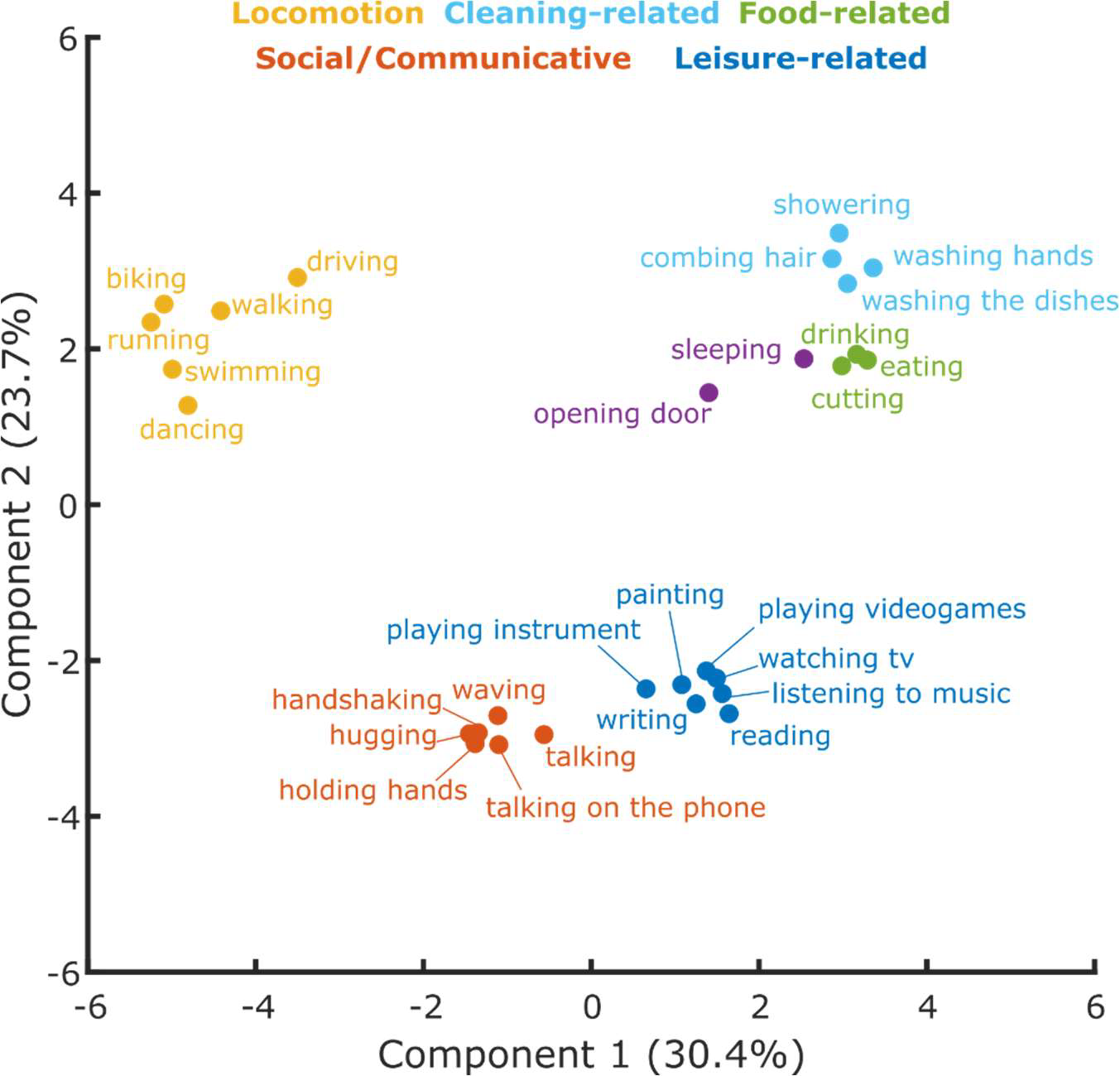
Cluster analysis. Clusters resulting from the K-means clustering analysis for the semantic model. The 2D-plot shows component 1 and 2, the corresponding labels of individual actions and the suggested labels for the categories resulting from the K-means clustering. For a visualization of the organization of these actions along the third component, which explained 23.4% of the variance, please refer to Figure S5 (Supplementary Information).

### RSA

To identify neural representations of observed actions that are organized according to semantic similarity, we performed a searchlight correlation-based RSA using the semantic model derived from the behavioral experiment. We thereby targeted brain regions in which the similarity of activation patterns associated with the observed actions matches the participants’ individual behavioral semantic similarity arrangement. We identified significant clusters in bilateral LOTC extending ventrally into inferior temporal cortex, bilateral posterior intraparietal sulcus (pIPS), and left inferior frontal gyrus/ventral premotor cortex (Figure 4, Table S2). The remaining models, focusing on action-related aspects, revealed some similarity with the semantic model (Figure 5A-D). This was expected since action-related features covary to some extend with semantic features (e.g. locomotion actions typically take place outdoors, cleaning-related actions involve certain objects, etc.; see also Figure S2). As explained above, an examination of the variance inflation factor indicated that these between-model correlations did not lead to problematic collinearities at the individual level, which allowed disentangling effects of the models using multiple regression. However, given these covariations, it is impossible to determine precisely what kind of information drove the RSA effects in the identified regions on the basis of the correlation-based RSA alone.

**Figure 4.**
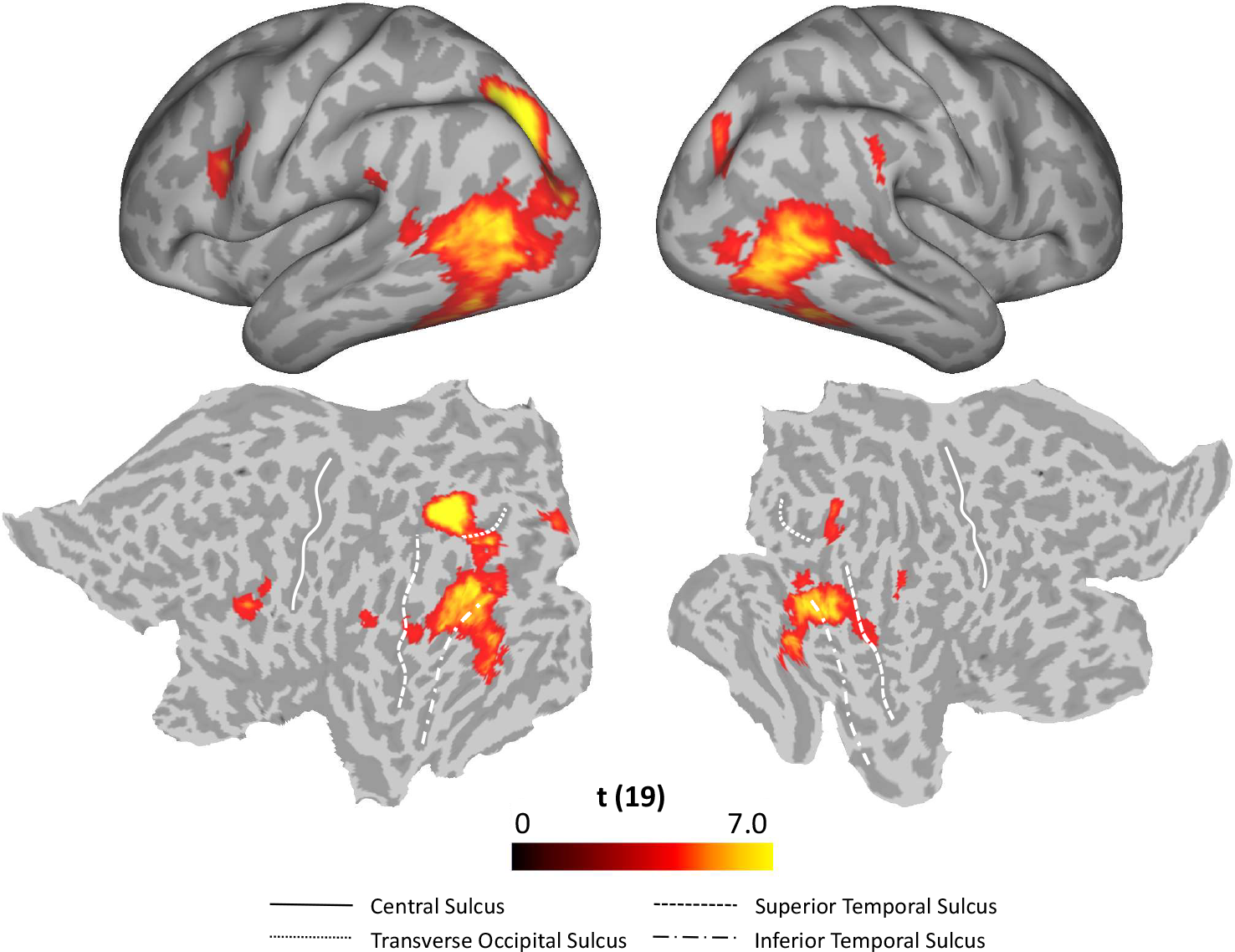
Standard RSA, Semantic model. Group results of the searchlight-based RSA using the semantic model (standard RSA, i.e. second order correlation between neural data and behavioral model). Statistical maps only show the positive t values that survived a cluster based nonparametric permutation analysis with Monte Carlo permutation (cluster stat: max sum; initial pval<0.001^16^). The resulting individual correlation maps were first fisher transformed and then converted to t scores. After the correction, data were converted to z scores, and only values greater than 1.65 (one-tailed test) were considered as significant. This analysis revealed clusters in bilateral LOTC, bilateral IPL and the left precentral gyrus (see Table S2 for details).

**Figure 5.**
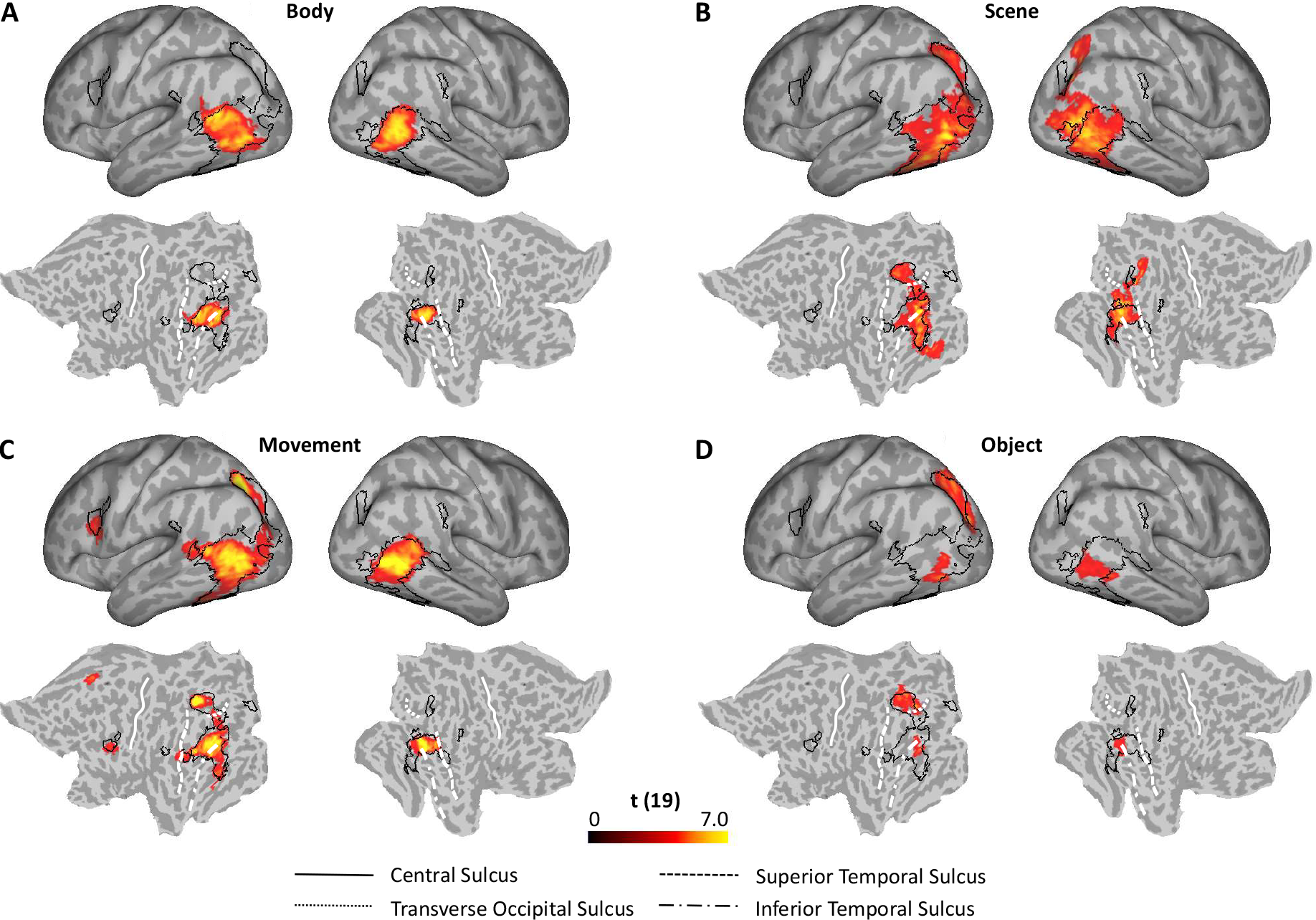
Standard RSA, control models compared to the semantic model. Statistical maps of the searchlight-based RSA for body (A), scene (B), movement (C), and object (D) models (see Table S2 for details). Black outlines indicate clusters of the correlation-based RSA for the semantic model (Figure 4). The analysis procedures and corrections were the same as for the semantic model (see Methods section and the caption of Figure 4).

Hence, to test which brain areas contain action information in their activity patterns that can be predicted by the semantic model over and above the action-related aspect models, we conducted a multiple regression RSA. We hypothesized that if actions were organized predominantly according to action-related aspects examined in the additional models (i.e., the body parts involved, the scene in which they typically occur, the movement kinematics, and the objects on which actions are performed), this analysis should not reveal any remaining clusters. As can be seen in Figure 6, the semantic model explained significant amounts of variance over and above the action-related aspect models in bilateral anterior LOTC at the junction to posterior middle temporal gyrus, right posterior superior temporal sulcus, and left pIPS.

**Figure 6.**
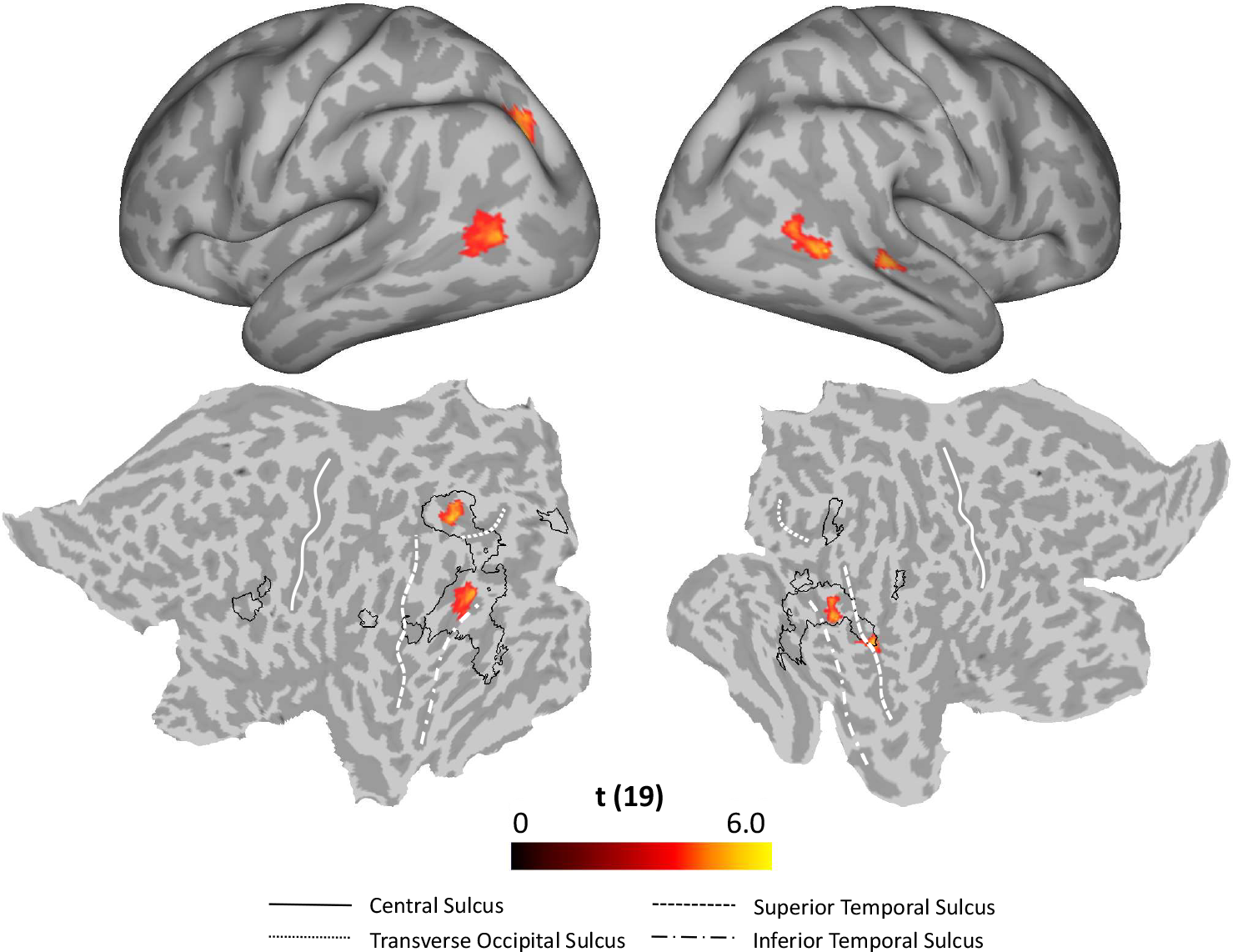
Multiple regression RSA, Semantic model. Group results of the searchlight-based multiple regression RSA, in which the five behavioral models were used as regressors in a multiple regression analysis conducted at the individual level. The resulting beta estimates were converted to t scores across participants and then corrected for multiple comparisons using cluster-based nonparametric permutation analysis^16^ (see Methods for details). Accounting for the remaining four models in the multiple regression analysis, the semantic model explained observed variance in bilateral LOTC and left IPL. For ease of comparison, the black outlines in the bottom part of the figure (flat maps) depict the clusters revealed by the standard RSA for the semantic model (Figure 4).

Figure S7 shows the results of the multiple regression RSA for the other models (body, scene, movement, and object models). For the body model (Figure S7A), we found clusters in left pSTS that explained significant amounts of variance over and above the remaining models. Likewise, we obtained a cluster in left dorsal middle occipital gyrus for the scene model (Figure S7B). The movement model explained observed variance over and above the other models in clusters partially overlapping with the clusters of the semantic model (Figure S7C). Notably, the clusters in LOTC found for the movement model were more posterior than those of the semantic model (for peak coordinates, see Table S3). By contrast, the cluster in pIPS were more anterior for the movement as compared to the semantic model. Finally, we obtained additional clusters in the left posterior fusiform gyrus and anterior middle temporal gyrus / superior temporal sulcus for the object model (Figure S7D).

Note that the obtained RSA results are unlikely to be due to some low-level features of the images we used. First, to minimize the risk that results could be driven by trivial pixel-wise perceptual similarities, we introduced a substantial amount of variability for each of the 28 actions, using different exemplars in which we varied the actor, the viewpoint, the background/ scene (e.g. kitchen A vs kitchen B), and the object (for actions that involved one). Second, a control analysis confirmed that there was no significant correlation between semantic and pixelwise similarity (see Results, Behavioral).

## Discussion

Here we aimed to investigate the organizational principles of everyday actions and the corresponding neural representations. Using inverse MDS^14^, we identified a number of clusters emerging in the arrangement of actions according to the meaning of actions that were relatively stable across participants. These clusters corresponded to meaningful categories, namely locomotion, communicative actions, food-related actions, cleaning-related actions, and leisure-related actions (Figure 3). Using multiple regression RSA^15^, we showed that this semantic similarity structure of observed actions was best captured in the patterns of activation in the LOTC (bilaterally) and left posterior IPL.

PCA suggested that the five categories revealed in the behavioral experiment appear to be organized along three main components that together explained around 78% of the observed variance. Whereas neither the PCA nor the k-means clustering provides objective labels of the main dimensions that might underlie the organization of actions into categories, it appears that clusters corresponding to semantic categories along the first principal component differed with respect to the type of change induced by the action (negative side of component 1: change of location, positive side of component 1: change of external/ physical state, middle: change of internal/ mental state). The second component seemed to distinguish actions that fulfil basic (or physiological) needs such as eating, drinking, cutting, or getting from one place to the other, and actions that fulfil higher (social belonging, self-fulfillment) needs such as hugging, talking to someone, reading, listening to music). Interestingly, this distinction shows some similarity with Maslow’s hierarchy structure of needs^17^. The third component might capture the degree to which an action is directed towards another person (hugging, holding hands, talking, etc.) or not (running, swimming, playing video games, reading). Note that we do not claim that the five categories obtained in the current study are the only action categories. We expect that future studies, using a similar approach as described in the current study with a wider range of actions, will reveal additional categories and overarching dimensions. However, given that we made an effort to select actions that we encounter on a regular basis, and given the agreement with semantic categories for words describing events, we can rule out that the categories obtained in the current study are entirely arbitrary.

To identify brain regions that encoded the similarity patterns revealed by the behavioral experiment, we conducted a searchlight-based RSA. We observed a significant correlation between the semantic model and the pattern of fMRI data in regions of the so-called action observation network^18^, which broadly includes the LOTC, IPL, premotor and inferior frontal cortex. Using multiple regression RSA, we found that only bilateral LOTC and the left angular gyrus/ pIPS contain action information as predicted by the semantic model over and above the remaining models. In line with these results, it has been demonstrated that it is possible to discriminate between observed actions based on the patterns of activation in the LOTC, generalizing across objects and kinematics^8^, and at different levels of abstraction^9^. Moreover, the LOTC has been shown to be sensitive to categorical action distinctions such as whether they are directed toward persons or objects^6^. Interestingly, studies using semantic tasks on actions using verbal stimuli^19^ or action classification across videos and written sentences^20^ tend to recruit anterior portions of the left LOTC, whereas studies using pictures find LOTC bilaterally and more posteriorly, closer to the cluster in the MOG identified in the current study. Together, these findings suggest that this area captures semantic aspects of actions at higher-order visual levels, whereas anterior portions of the left LOTC might capture these aspects at stimulus-independent or verbal levels of representation (see also^21^).

Another novel finding of our study was that semantic similarity of actions is also encoded in the left angular gyrus/pIPS. Interestingly, an area posteriorly/ventrally of pIPS at the dorsal part of left middle occipital gyrus/retinotopic area V3A & V3B (V3CD), posterior to the transverse occipital sulcus (TOS)^22^, also named occipital place area (OPA)^22,23^, was found to represent the similarity of scenes in which the actions took place. OPA has been shown to respond to local elements of scenes (e.g. furniture, floor) rather than global scene properties (such as boundaries and spatial expanse, i.e., the gist of a scene)^24^. Moreover, OPA has been demonstrated to be sensitive to the navigational affordances of a scene (i.e., the pathways for movements one may use to navigate through the scene)^25^. This profile is not only important for visually-guided navigation and obstacle avoidance, but may also provide critical cues indicating potential interactions with the local environment. Being part of the dorsal “where” pathway, the IPS is known to be sensitive to spatial aspects of actions. Thus, a possible interpretation is that pIPS captures relationships between an action and the environment/scene in which an action takes place. In analogy to *object affordances*, we suggest to refer to such scene-based constraints of action possibilities as *scene affordances*^26^ or *scene functions*^27^. The anatomical dissociation between scene-related and semantic effects in MOG/V3CD and pIPS, respectively, might point to distinct but functionally related specializations for scene-related and action-related information, respectively. In line with this view, MOG/TOS shows enhanced neural responses for the sight of manipulable, contextual objects in the periphery of an action, whereas IPS is modulated by the goal-relatedness of contextual objects with an action^28^. This suggests that MOG/V3CD registers local, potentially action-relevant elements in a scene, whereas pIPS integrates this information with semantic aspects of the observed action. Interestingly, previous MVPA studies did not associate pIPS with the representation of action information. So far, no study investigated the representational organization of actions that were embedded in different contextual scenes. Hence, scene-specific aspects of action, such as the functional relationship between actions and scenes (e.g. cooking-kitchen, running-park, etc.), could not be captured by previous studies.

Our results call for a comparison with the object domain, where similar questions have been addressed for decades. In line with the results of the GLM-based RSA, both the pIPS and the LOTC have been demonstrated to represent the similarity structure of object categories^29^. Regarding the results of the cluster analysis, salient distinctions at the behavioral and neural level are found between animate and inanimate objects^30,31,e.g. 32^, which are further segregated into human and nonhuman objects^33^, and manipulable and non-manipulative objects^34^, respectively. The division between animate and inanimate objects, supported by neuropsychological, behavioral and neuroimaging findings, has been suggested to have a special status, likely due to evolutionary pressures that favored fast and accurate recognition of animals^32,33,35^. We conjecture that similar evolutionary mechanisms might have produced the distinction between actions belonging to different categories, such as locomotion (which might indicate the approach of an enemy), food-related actions (which might be critical for survival) and communicative actions (critical for survival within a group).

Finally, our data showed that people tended to organize actions based on dissimilarity regarding their meaning and the scenes in which they take place in a similar way (Figure 3 and Figure S6B). This observation is corroborated by studies investigating scene categorization: the way humans categorize scenes can be predicted best by a model that describes which actions can be carried out in a scene^27^. In line with our findings, this model is also capable to predict the representational organization of observed scenes in left LOTC, even if scenes only rarely contained persons or salient objects that are associated with specific actions^36^. These results support the idea that there might be a strong interplay between the neural mechanisms involved in processing the scene and the meaning of an action, most likely due to the statistical co-occurrence of certain actions taking place in some scenes (such as food-related actions and kitchen) in comparison to others (such as food-related actions and a garage).

## Conclusions

Using a combination of behavioral and fMRI data, we identified a number of meaningful semantic categories according to which participants arrange observed actions. The corresponding similarity structure was captured in the LOTC and the pIPS over and above the major elements of perceived actions (body parts, scenes, movements, and objects), supporting the view that these areas play a critical role in accessing the meaning of actions beyond the mere perceptual processing of action-relevant features. We propose that the LOTC encodes conceptual aspects of actions (e.g. that both riding a bike and swimming can be considered a form of locomotion) whereas pIPS might contribute towards the representation of action semantics by extracting and combining relevant elements of action scenes (e.g. the objects in a scene, the posture and orientation of the person, the underlying movement kinematics).

## Methods

### Participants

Twenty healthy participants (13 females; mean age: 28 years; age range: 20 – 46) took part in an fMRI and a behavioral experiment carried out at the Combined Universities Brain Imaging Centre (CUBIC) at Royal Holloway University of London (RHUL). The experiment was approved by the RHUL ethics committee. All participants were right-handed with normal or corrected-to-normal vision and no story of neurological or psychiatric disease. Participants received and gave written informed consent before starting the experiment.

### Inverse Multidimensional Scaling

Participants sat in front of a monitor (LCD 16.2×19.2 inches; distance 60 cm). In trial 1, all action images (1 exemplar per action) appeared on the screen in a circular arrangement (with the order of actions randomly selected; see Figure 1B). Participants were instructed to arrange the pictures by drag-and-drop using the mouse according to their perceived similarity in meaning (e.g. *walking* and *running* would be placed closer to each other than *walking* and *drinking*) and to press a button once they were satisfied with the arrangement. In each subsequent trial (trial 2 to N_p_, where N_p_ is the total number of trials for participant *p*), a subset of stimuli was sampled from the original stimulus set. The subset of actions was defined using an adaptive algorithm that provided the optimal evidence for the pairwise dissimilarity estimates (which are inferred from the 2D arrangement of the items on the screen, see^14^ for details).

### Stimulus selection

In contrast to previous studies that used a small set of actions, we aimed to cover a wide range of actions that we encounter on a daily basis. To this aim, we initially carried out an online survey using Google Forms. The aim of the survey was to identify actions that are considered common by a large sample of people. We thus asked 36 participants (different from those that took part in the fMRI study) to spontaneously write down all the actions that came to their mind within 10 minutes that they or other people are likely to do or observe. As expected, participants often used different words to refer to similar meanings (e.g. talking and discussing) or used different specific objects associated with the same action (e.g. drinking coffee and drinking water). Two of the authors (EB, AL) thus assigned these different instances of similar actions to a unique label. Actions were selected if they were mentioned by at least 20% of the participants. In total, we identified 37 actions (see Table S1).

As a next step, we selected a subset of actions that were best suitable for the fMRI experiment. Specifically, we aimed to choose a set of actions that were arranged consistently across participants according to their perceived similarities in meaning. To this aim, we retrieved images depicting the 37 actions from the Internet. Using these images, we carried out inverse MDS (see corresponding section and ^14^ for details) using 15 new participants. Each participant had 20 minutes to complete the arrangement. In three additional 20-minute sessions, participants were furthermore instructed to arrange the actions according to the perceived similarity in terms of the scenes in which these actions typically take place, movement kinematics, and the objects which are typically associated with these actions. The order in which these four tasks (semantics, scenes, movements, objects) were administered to participants were counterbalanced across participants. To rule out that any obtained arrangements were driven by the specific exemplars chosen for each action, we repeated the same experiment with a new group of people (N=15) and an independent set of 37 images taken from the Internet.

To construct representational dissimilarity matrices (RDMs), we averaged the dissimilarity estimates for each pair of actions (e.g. the dissimilarity between biking and brushing teeth, etc.), separately for each participant and each task, across trials. For each participant, we then constructed dissimilarity matrices based on the Euclidean distance between each pair of actions that resulted from the inverse MDS experiment. The dissimilarity matrices were then normalized by dividing each value by the max. value of each matrix. Each row of this matrix represented the dissimilarity judgment of one action with respect to every other action. To select the most suitable actions for the fMRI experiment, we aimed to evaluate which of the 37 actions were arranged similarly across participants in the different tasks. To this aim, we carried out a cosine distance analysis, which allowed us to determine, for each action, the similarity across all participants. The cosine distance evaluates the similarity of orientation between two vectors. The cosine distance can be defined as one minus the cosine of the angle between two vectors of an inner product space: a cosine distance of 1 indicates that the two vectors are orthogonal to each other (maximum dissimilarity/minimum similarity); a cosine distance of zero indicates that the two vectors have the same orientation (maximum similarity/minimum dissimilarity). The cosine distance can therefore range between 0 and 1. In an RDM, each row (or column) represents the dissimilarity score between one action and every other action, ranging from 0 (minimum dissimilarity) to 1 (maximum dissimilarity). Therefore, each row of the matrix of each single participant was used to compute the pairwise cosine distances between this and the corresponding row of every other participant. For each action, a small cosine distance between two single participants would indicate that they agreed on the geometrical configuration of that action with respect to every other action; a value close to 1 would indicate disagreement. For each action, we computed the mean across the pairwise cosine distances of all participants in both behavioral pilot experiments and kept only those actions that had a cosine distance within one standard deviation from the averaged cosine distance in all tasks and both stimulus sets. Thirty-one actions fulfilled this criterion, whereas five (*getting dressed, cleaning floor, brushing teeth, singing and watering plants*) had to be discarded. We also decided to remove two additional actions (*grocery shopping and taking the train*) because these could not be considered as single actions but implied a sequence of actions (e.g. *entering the shop, choosing between products, etc.; waiting for the train, getting on the train, sitting on the train, etc*.).

At the end of the procedure, we identified 30 actions that could be used for the next step which consisted in creating the final stimulus dataset. To this aim, we took photos of 29 of the 30 actions using a Canon EOS 400D camera. To maximize perceptual variability within each action, and thus to minimize low-level feature differences between actions, we varied the actors (2), the scene (2) and perspectives (3), for a total of 12 exemplars per action. Exemplars for the action ‘swimming’ were collected from the Internet because of the difficulties in taking photos in a public swimming pool.

The distance between the camera and the actor was kept constant within each action (across exemplars). Since some actions consisted of hand-object interactions (such as painting, drinking) and thus required finer details while other actions involved the whole body (such as dancing, running) and thus required a certain minimum distance to be depicted, it was not possible to maintain the same distance across all the actions. The two actors were instructed to maintain a neutral facial expression and were always dressed in similar neutral clothing. If an action involved an object, the actor used two different exemplars of the object (e.g. two different bikes for *biking*) or two different objects (e.g. a sandwich or an apple for *eating*). Furthermore, some actions required the presence of an additional actor (*handshaking, hugging, talking*). The brightness of all pictures was adjusted using PhotoPad Image Editor (www.nchsoftware.com/photoeditor/). Pictures were then converted into greyscale and resized to 300 × 400 pixels using FastOne Photo (www.faststone.org). In addition, we made the images equally bright using custom written Matlab code (mean brightness across all images was 115.80 with standard deviation equal to 0.4723).

To ensure that the final set of pictures were comprehensible and identified as the actions we intended to investigate, we furthermore validated the stimuli through an online survey using Qualtrics and Amazon Mechanical Turk using 30 participants. Specifically, the 30*12 = 360 pictures were randomly assigned to three groups of 120 images. Each group was assigned to ten participants that had to name the actions depicted in the images. For each participant, the images were presented in a random sequence. Since most of the participants failed to correctly name some of the exemplars of *making coffee* and *switching on lights*, these actions were excluded from the stimulus set. Therefore, the final number of actions chosen for the main experiment was 28 (see Table S1).

### Experimental design and setup

The fMRI experiment consisted of twelve functional runs and one anatomical sequence halfway through the experiment. Each functional run started and ended with 15 seconds of fixation. In between runs, the participants could rest.

Stimuli were back-projected onto a screen (60Hz frame rate) via a projector (Sanyo, PLC-XP-100L) and viewed through a mirror mounted on the head coil (distance between mirror and eyes: about 12 cm). The background of the screen was uniform grey. Stimulus presentation and response collection was controlled using ASF^37^, a toolbox based on the Matlab Psychophysics toolbox^38^.

Each functional run consisted of 56 experimental trials (28 exemplars of the action categories performed by each of the two actors) and 18 null events (to enhance design efficiency) presented in a pseudorandomized order (preventing that the same action was shown in two consecutive trials, except during catch trials, see next paragraph). A trial consisted of the presentation of an action image for 1 second followed by 3 s of fixation. A null event consisted of the presentation of a fixation cross for 4 s. Within run 1-6, each possible combination of action types (28) × exemplars (12) was presented once, for a total of 336 trials. For runs 7-12, the randomization procedure was repeated such that each possible combination was presented another time. In this way, each participant saw each exemplar twice during the entire experiment (and thus each action was presented 24 times). A full balancing of all combinations of action, scene, and actor within each run was not possible with 28 actions, therefore the experiment was quasi-balanced: in each run, if actor 1 performed an action in scene A, actor 2 performed the same action in scene B, and vice versa.

To ensure that participants paid attention to the stimuli, we included seven (out of 63; 4.41%) catch trials in each run which consisted in the presentation of an image depicting the same action (but not the same exemplar) as the action presented in trial N-1 (e.g. *eating* an apple, actor A, scene A, followed by *eating* a sandwich, actor B, scene B). Participants were instructed to press a button with their right index finger whenever they detected an action repetition. Within the entire experimental session, all 28 actions could serve as catch trials and each action was selected randomly without replacement such that the same action could not be used as a catch trial within the same run. After a set of 4 runs all 28 actions were used as catch trial once, thus the selection process started from scratch. Catch trials were discarded from multivariate data analysis.

Before entering the scanner, participants received written instructions about their task and familiarized with the stimulus material for a couple of minutes. Next, participants carried out a short practice run to ensure that they properly understood the task.

### MRI data acquisition

Functional and structural data were acquired using a Siemens TIM Trio 3T MRI scanner. For the acquisition of the functional images, we used a T2*-weighted gradient EPI sequence. The repetition time (TR) was 2.5 seconds, the echo time (TE) was 30 milliseconds, the flip angle was 85°, the field of view was 192 × 192 mm, the matrix size was 64 × 64, and the voxel resolution was 3 × 3 × 3 mm. A total of 37 slices were acquired in ascending interleaved order. Each functional run lasted 5 minutes and 55 seconds and consisted of 142 volumes.

For the structural data, we used a T1*-weighted MPRAGE sequence (image size 256 × 256 × 176 voxels, voxel size 1 × 1 × 1 mm, TR 1.9 sec, TE 3.03, flip angle 11), lasting 5 minutes and 35 seconds.

### MRI data preprocessing

Anatomical data were segmented using FreeSurfer^39^. Preprocessing of the functional data was carried out using SPM12 (http://www.fil.ion.ucl.ac.uk/spm/software/spm12/). The slices of each functional volume were slice time corrected and then spatially realigned to correct for head movements. Functional volumes were then coregistered to the individual anatomical image. Analyses were conducted in individual volume space, but using the inner and outer surfaces obtained with FreeSurfer as a constraint to select the voxels included in each searchlight as implemented in CoSMoMVPA^40,41^. The resulting maps were resampled to the surface level on the Human Connectome Project common space *FS_LR 32k*^42^ using FreeSurfer and workbench connectome^43^. Multivariate analyses were conducted using unsmoothed data.

### Behavioral experiment and model selection

Following the fMRI experiment, either on the same or the next day, participants took part in an additional behavioral experiment in which they carried out an inverse MDS task using similar procedures as described above (see **Methods** section and Figure 1B), using the same actions that were used during the fMRI experiment. In separate blocks of the experiment, participants were asked to arrange the actions according to their perceived similarity in terms of (a) meaning, (b) the body part(s) involved, (c) scene, (d) movement kinematics, (e) the object involved. The order of blocks was counterbalanced across participants.

To construct dissimilarity matrices for each participant and task (a – e), we used the same procedure described in the section **Stimulus selection**, i.e. we determined the Euclidean distance between each pair of actions that resulted from inverse MDS, and normalized the dissimilarity matrices by dividing each value by the maximum value of each matrix. Individual dissimilarity matrices were used as a model for the representational similarity analysis of fMRI data (see next section).

To further characterize the structure that emerged from the inverse MDS, we adopted principal component analysis (PCA) as implemented in the R package *cluster* to individuate the principal components along which the actions were organized. To characterize the observed clusters, we furthermore used a model-based approach using the *K-means*^44^ clustering method. The K-means method requires the number of clusters as an input, which was one of the parameters we wished to estimate from the data. To this aim, we used the Silhouette method^45^ as implemented in the R package *factoextra* to estimate the optimal number of clusters. Specifically, this method provides an estimate of the averaged distance between clusters as a function of the number of clusters used and selects the value which provides the maximal distance.

### Representational Similarity Analysis (RSA)

The aim of the RSA was to individuate those brain regions that were best explained by the models obtained behaviorally and thus to infer the representational geometry that these areas encoded. We therefore conducted an RSA over the entire cortical surface using a searchlight approach^46^ at the individual brain space. Each searchlight consisted of 100 features (1 central vertex + 99 neighbors). All multivariate analyses were carried out using custom written Matlab functions and CoSMoMVPA^41^.

For the multivariate analysis, the design matrix consisted of 142 volumes X 28 predictors of interest (resulting from the 28 actions) plus nuisance predictors consisting of the catch trials, the parameters resulting from motion correction, and a constant term. Thus, for each participant, we obtained 28 beta maps at the volume level. The beta maps were averaged across runs and normalized (z transformed) across voxels of each searchlight before the analysis. We adopted two approaches for the representational similarity analysis, a standard and a multiple regression RSA approach.

#### Standard RSA

First, to identify clusters in which the neural data reflected the dissimilarity patterns derived from the different behavioral tasks, we conducted a standard RSA in which we correlated the dissimilarity matrix (DSM) derived from the neural data of each searchlight with the normalized dissimilarity matrices (DSMs) of each of the five behavioral models. The correlation values were assigned to the central node of each searchlight, thus leading to a correlation map for each of the five models, separately for each participant. The correlation maps of this first-level analysis were then resampled to the common space and fisher transformed to normalize the distribution across participants to run a second-level analysis. Specifically, for each of the five models, the correlation maps of all N participants were tested against zero using a t-test at each vertex. The resulting t maps were corrected using a cluster-based nonparametric Monte Carlo permutation analysis (5000 iterations; initial threshold p < 0.001^16^).

#### Multiple regression RSA

To determine clusters in which the neural DSM correlated with a given behavioral DSM while accounting for the remaining behavioral DSMs, we conducted a multiple-regression analysis at each searchlight in which we used the DSMs derived from the five models as regressors of interest to evaluate which predictor best explained the observed variance. This analysis provided us with beta maps for each predictor (i.e., behavioral model) that were then entered in a second-level (group) analysis to test the individual beta maps against zero. The procedure for multiple comparisons correction was the same as described in the previous paragraph.

To estimate potential risks of collinearity, we computed the Variance Inflation Factor (VIF) for each participant to have a measure of the *inflation* of the estimated variance of the *ith* regression coefficient (computed as 1/(1−R^2^_i_), where *i* indicates a variable and R^2^ is the coefficient of determination), assuming this coefficient being independent from the others. The VIFs were relatively small (average VIF semantic model: 1.52, body model: 1.29, scene model: 1.51, movement model: 1.29, object model: 1.28), indicating a low risk of multicollinearity^47^ and thus justifying the use of multiple regression analysis.

Behavioral and fMRI data, materials and code used in the current study are available from the corresponding author upon request.

## Acknowledgements

This work was supported by Royal Holloway University of London and a grant awarded to Angelika Lingnau (Heisenberg-Professorship, German Research Foundation, Li 2840/2-1).

## Supplementary Information

**Figure S1.**
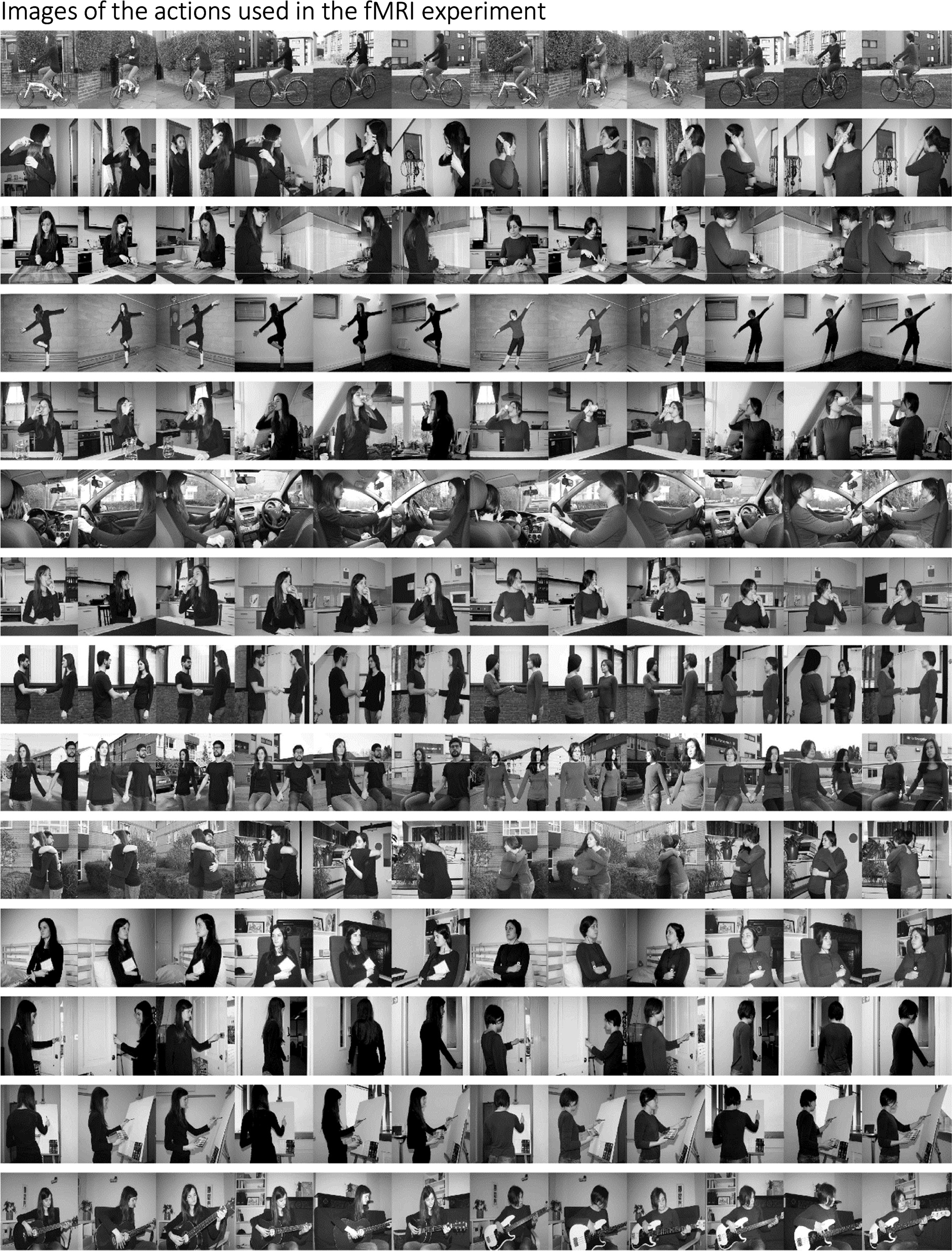
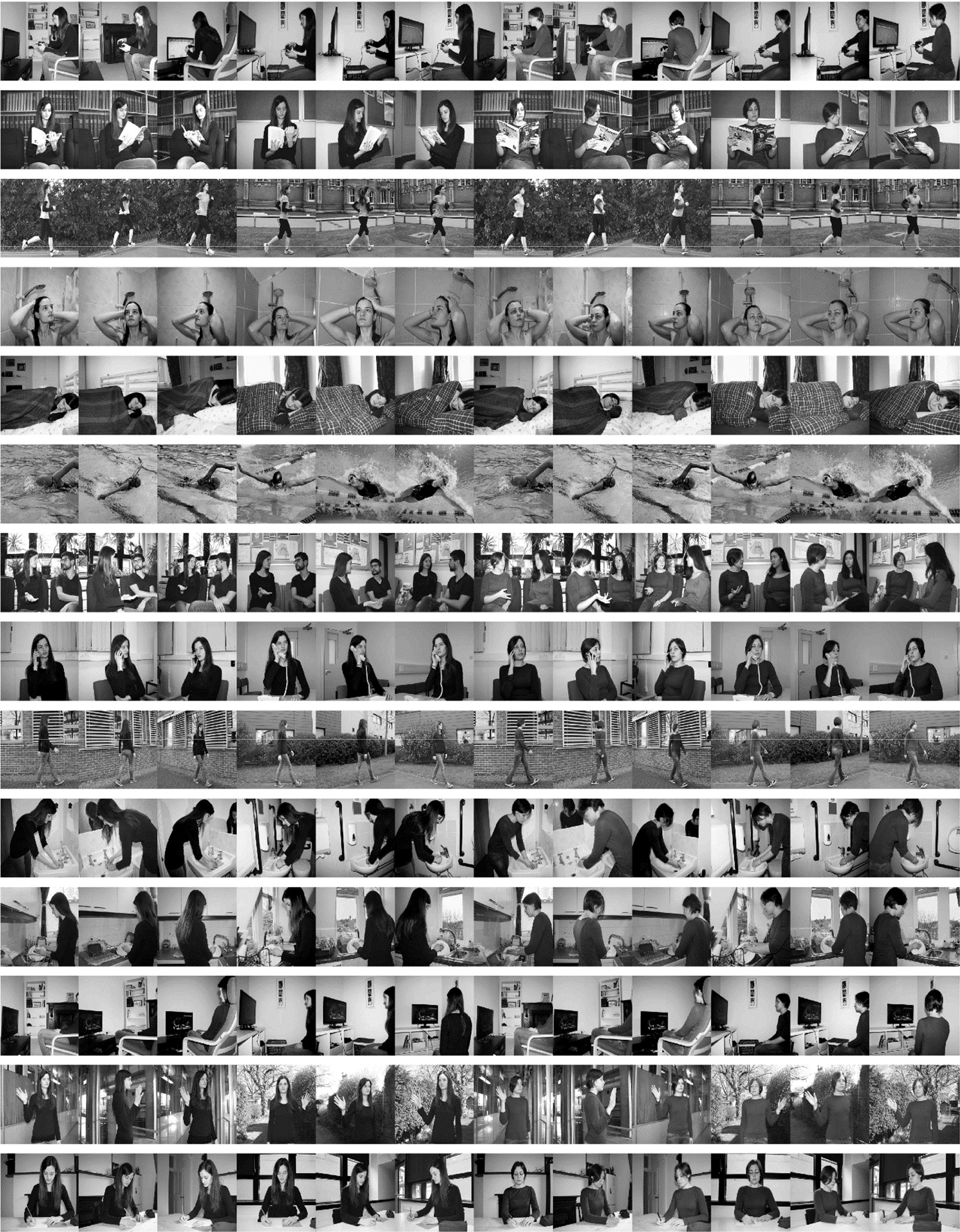
All images used in the fMRI experiment. Different actions are depicted in rows (for corresponding labels, see words printed in bold font in Table S1). Different exemplars for each action are shown in the columns. Exemplars varied in terms of the actor (2), scene (2), and viewing angle (3).

**Figure S2.**
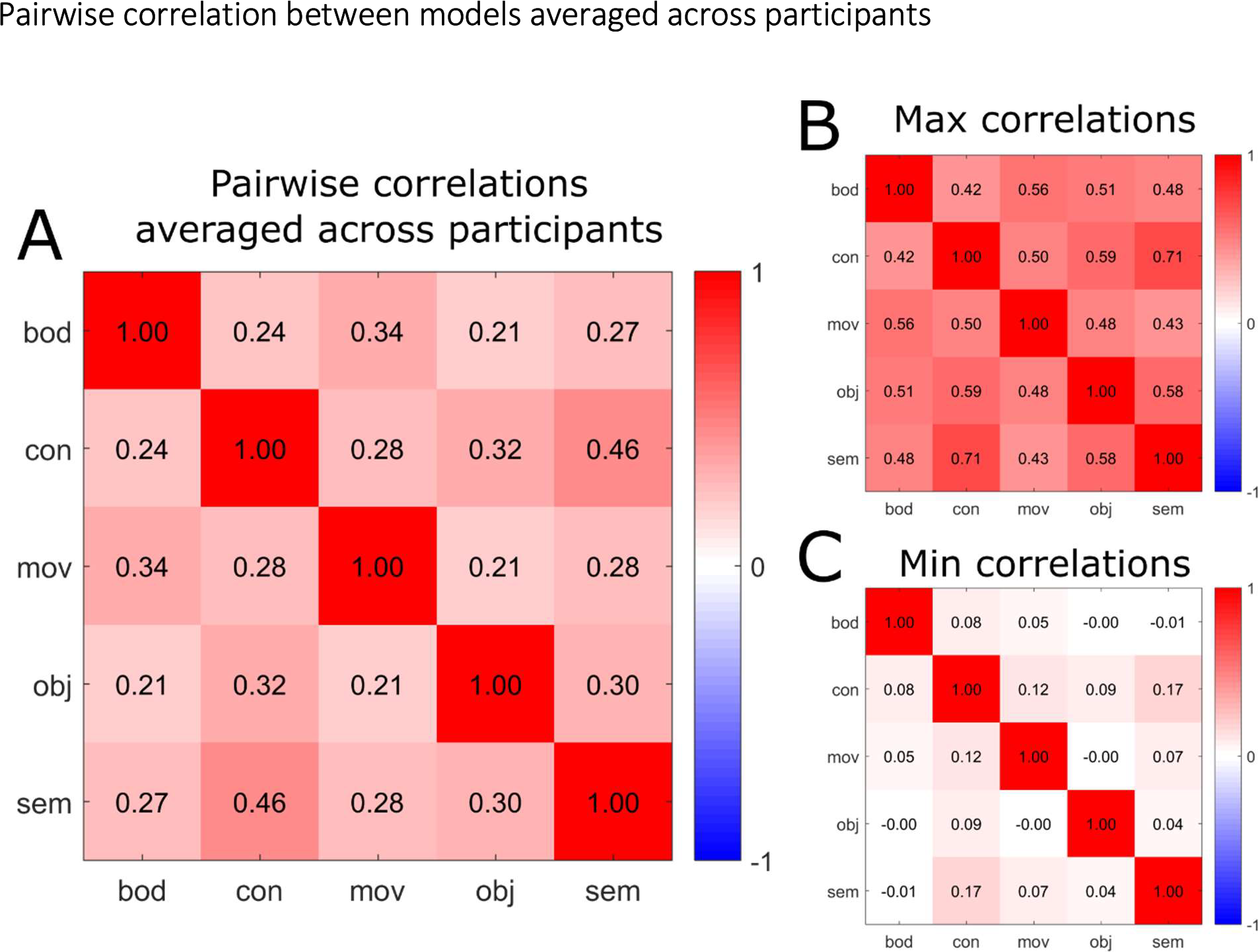
Pairwise cross-correlation matrix across models. For each participant, we computed the pairwise cross-correlation matrix across models. We then computed (A) the averaged correlation values across participants; and reporting (B) the maximum and (C) the minimum correlation values across participants. For a corresponding analysis of collinearity using the variance inflation factor, see Methods, Multiple Regression RSA.

**Figure S3.**
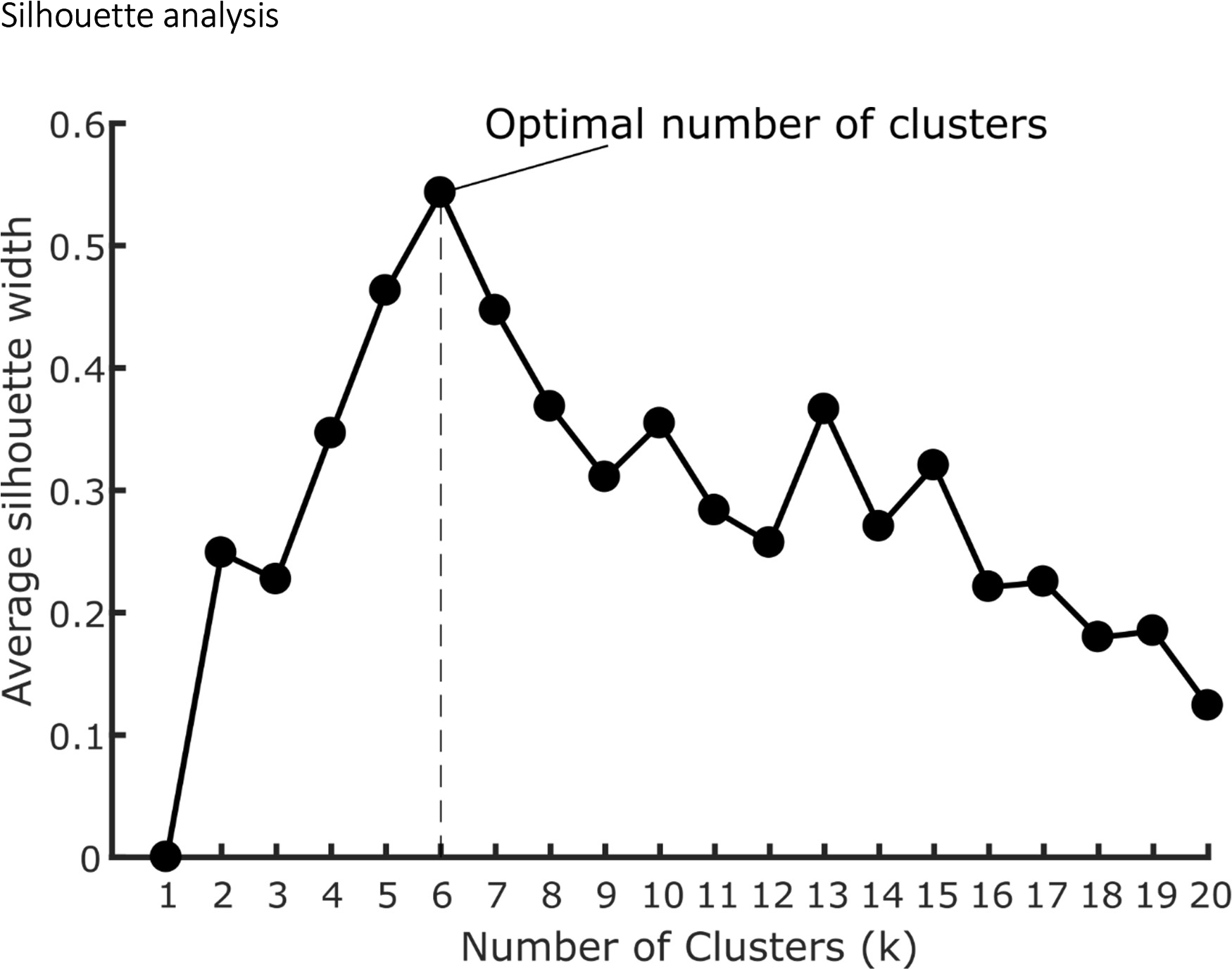
Silhouette analysis revealed that the optimal number of clusters for the Semantic model was 6. The analysis was performed using the fviz_nbclust function of the R package “factoextra”. The silhouette width is a measure of how well each item fits within its cluster with higher values indicating better fit. The average silhouette width is the average of the all items. The algorithm computes the average silhouette width as a function of the number of clusters (k). The highest value of the average silhouette width is taken and the corresponding k is selected as the optimal number of clusters.

**Figure S4.**
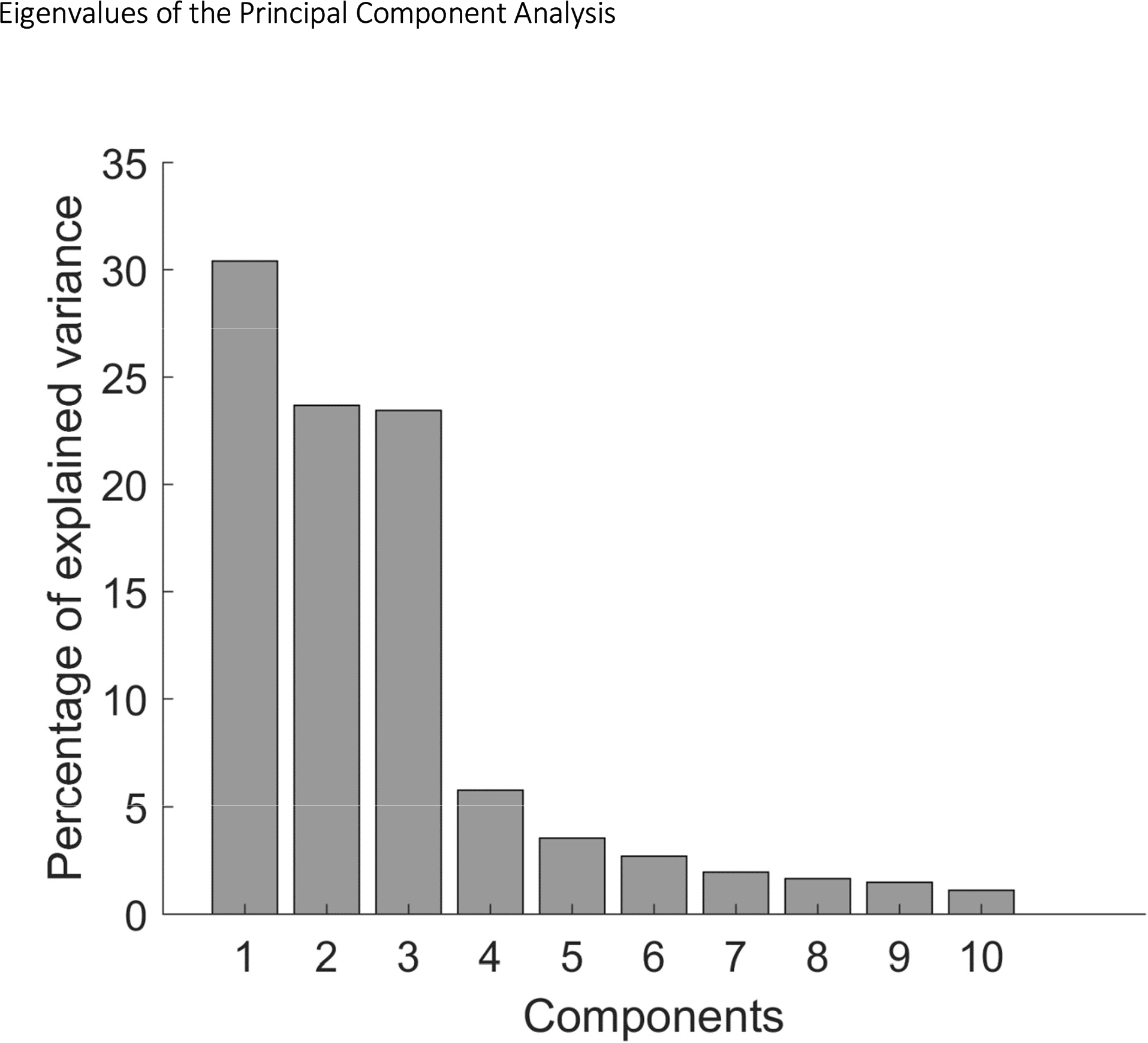
Eigenvalues obtained from the PCA of the semantic model. The PCA shows that the first 3 components accounted for the largest amount of variance

**Figure S5.**
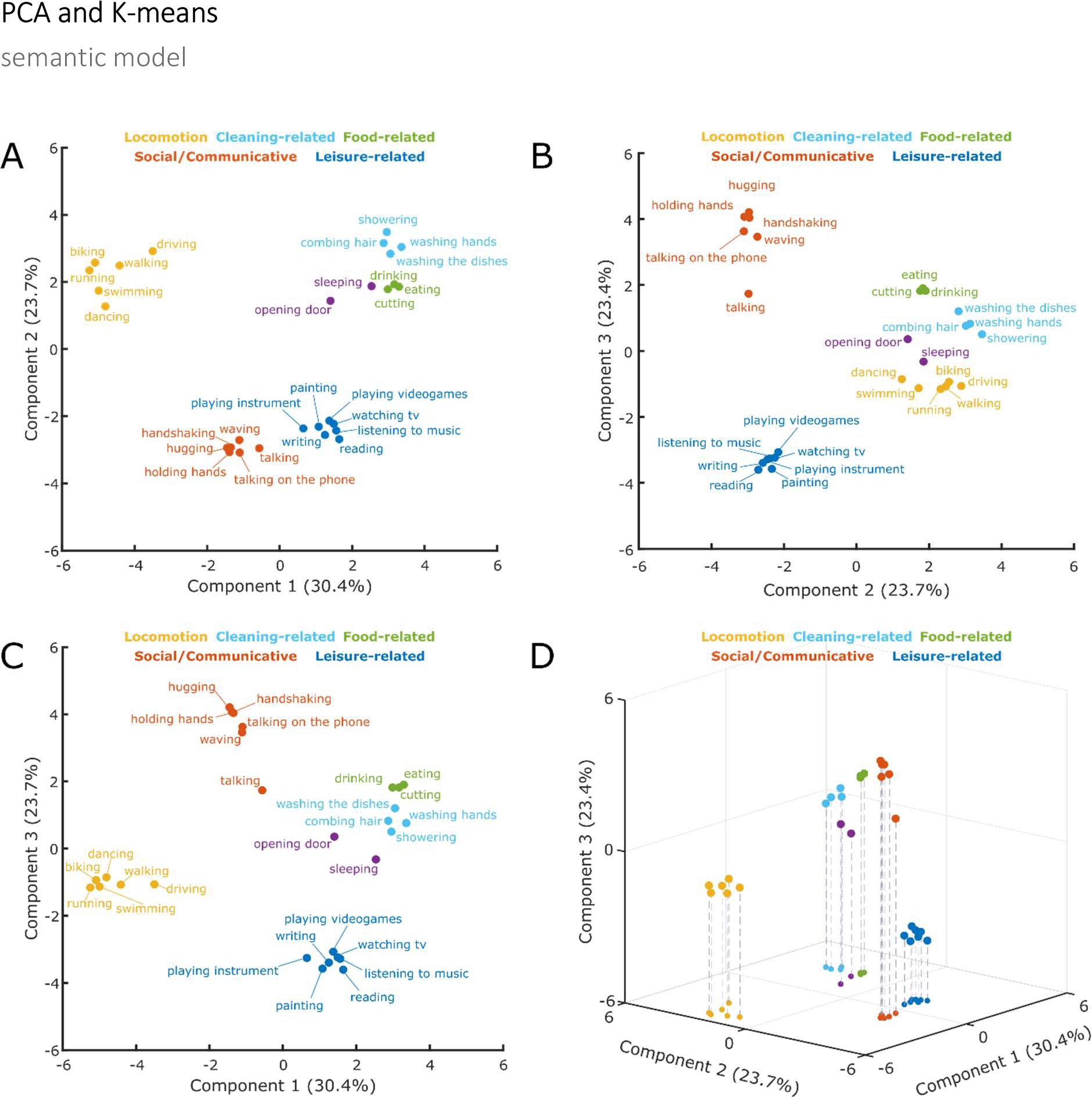
Cluster analysis. Clusters resulting from the K-means clustering analysis (K-means) for the semantic model. A: 2D-plot showing component 1 and 2, corresponding labels of individual actions and suggested labels for the categories resulting from the K-means clustering. B: 2D-plot showing component 2 and 3. C: 2D-plot showing component 1 and 3. D: 3D-plot showing components 1-3.

**Figure S6.**
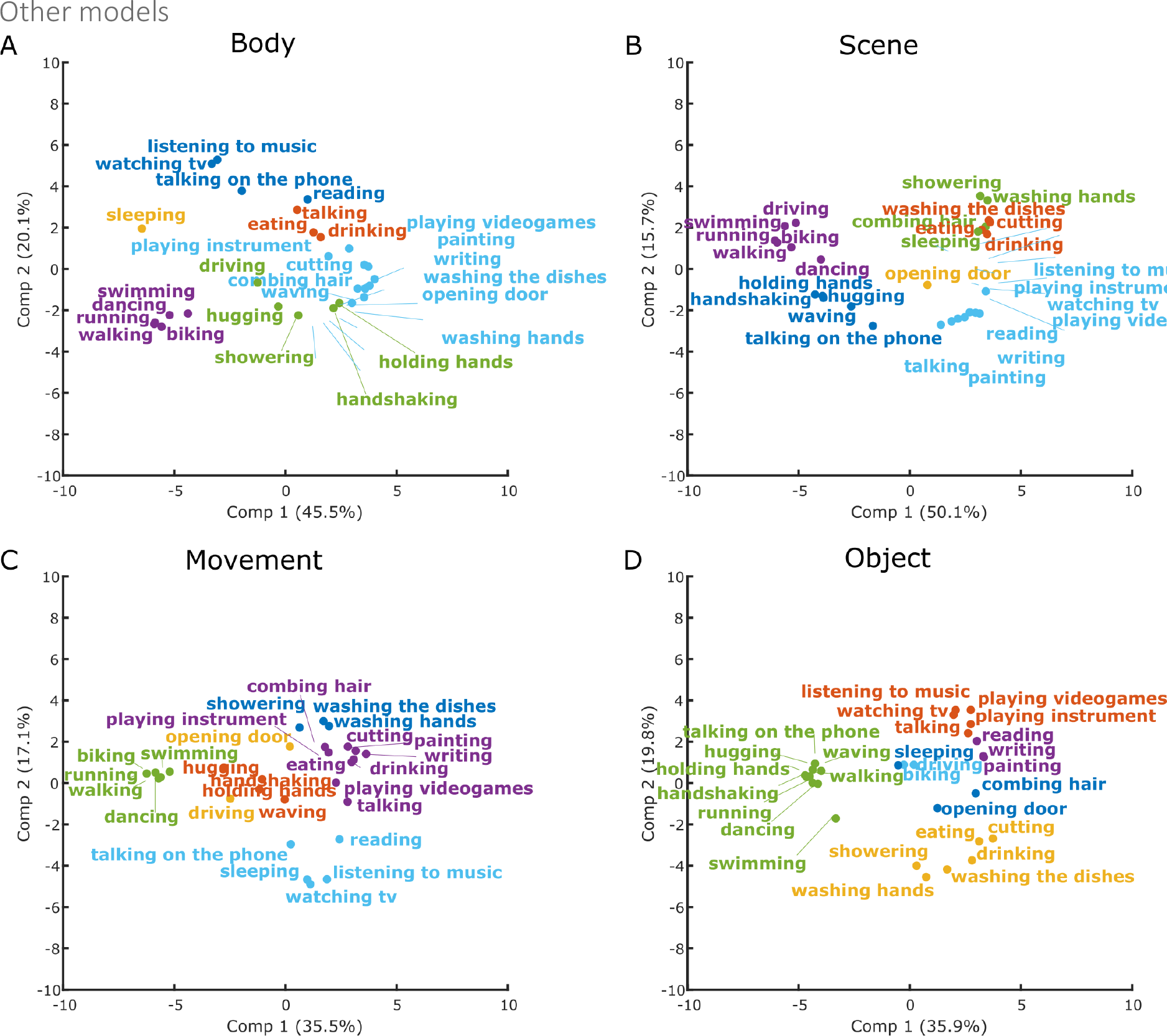
First 2 principal components of the control models. Clusters were distinguished using the K-Means methods as implemented in R. Number of clusters used as input were 6 (based on the Silhouette analyses of the semantic model; see Figure S3). Colors are used to distinguish between the six resulting clusters in each analysis.

**Figure S7.**
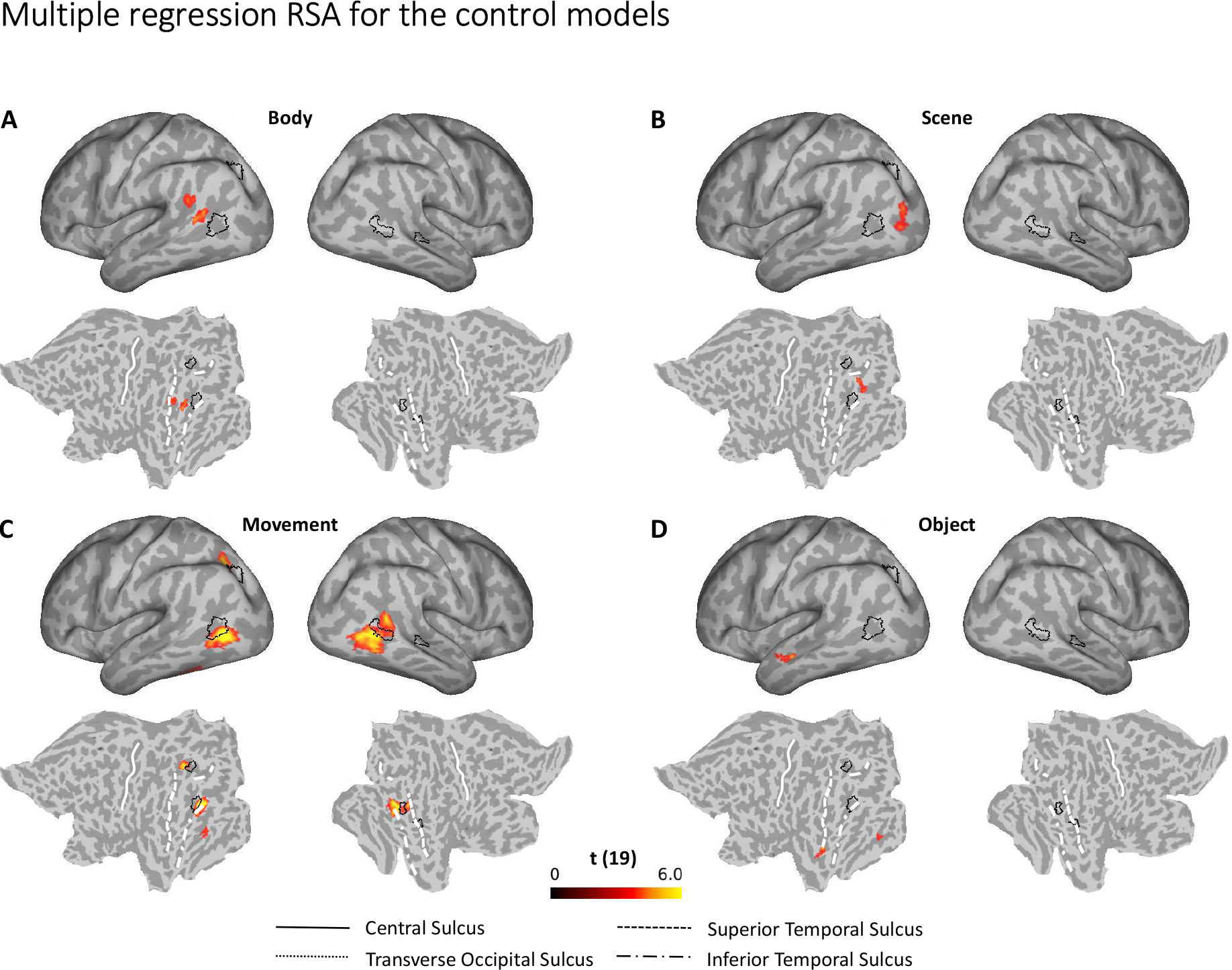
Multiple regression RSA, comparison between the semantic and the other models. Group results of the searchlight-based multiple regression RSA for the body (A), scene (B), movement (C), and object (D) model. For details, see methods section and captions of Figure 4. Black outlines on the inflated brains and the flat maps depict significant clusters revealed by the multiple regression RSA for the semantic mode (Figure 6).

**Table S1.**
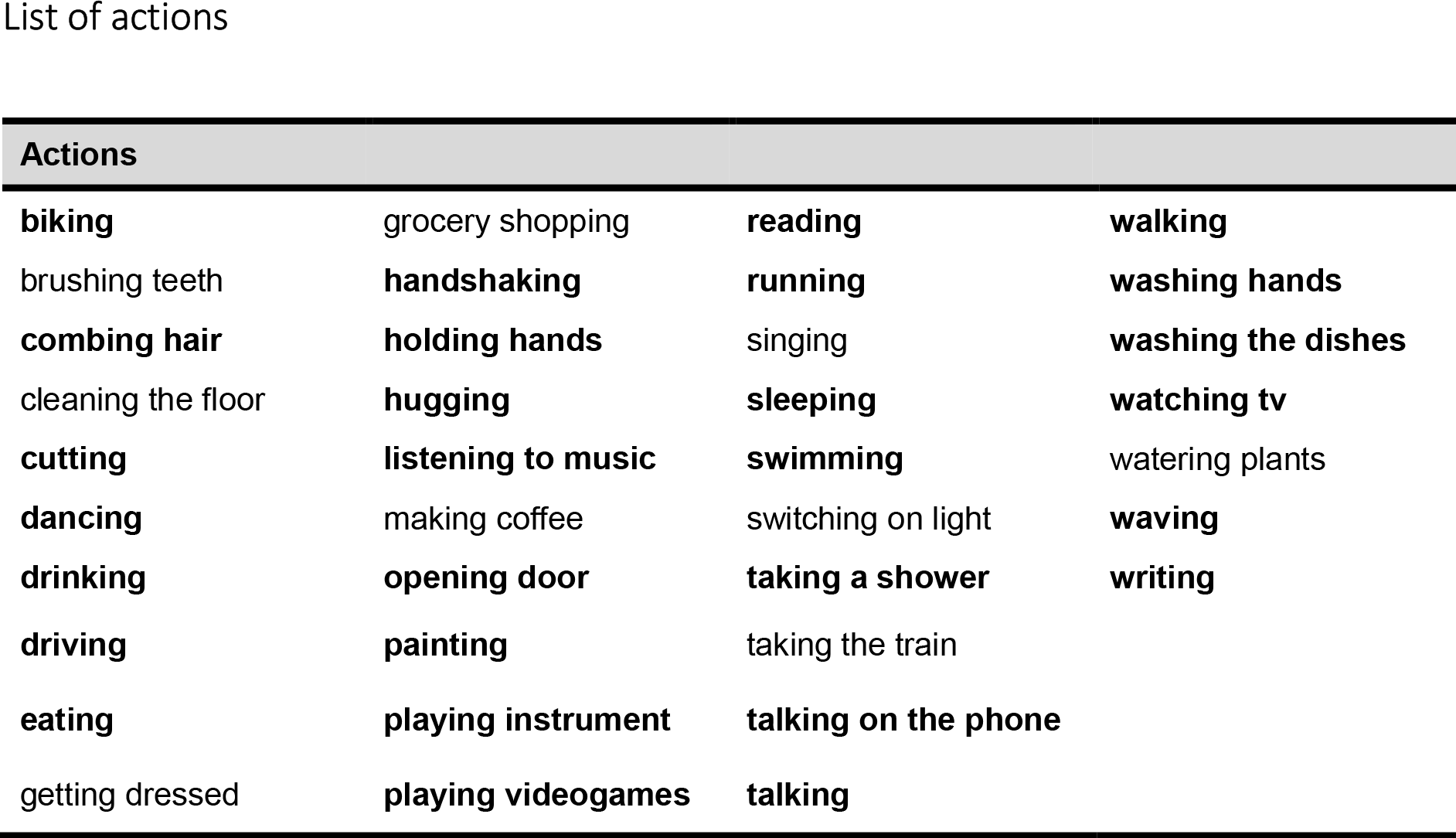
List of actions identified using the online survey. We initially kept actions that were mentioned by at least 20% of the participants. The final selected twenty-eight actions used for the study are the ones highlighted. See the Methods section of the main article for the procedure we used to select these actions.

**Table S2.**
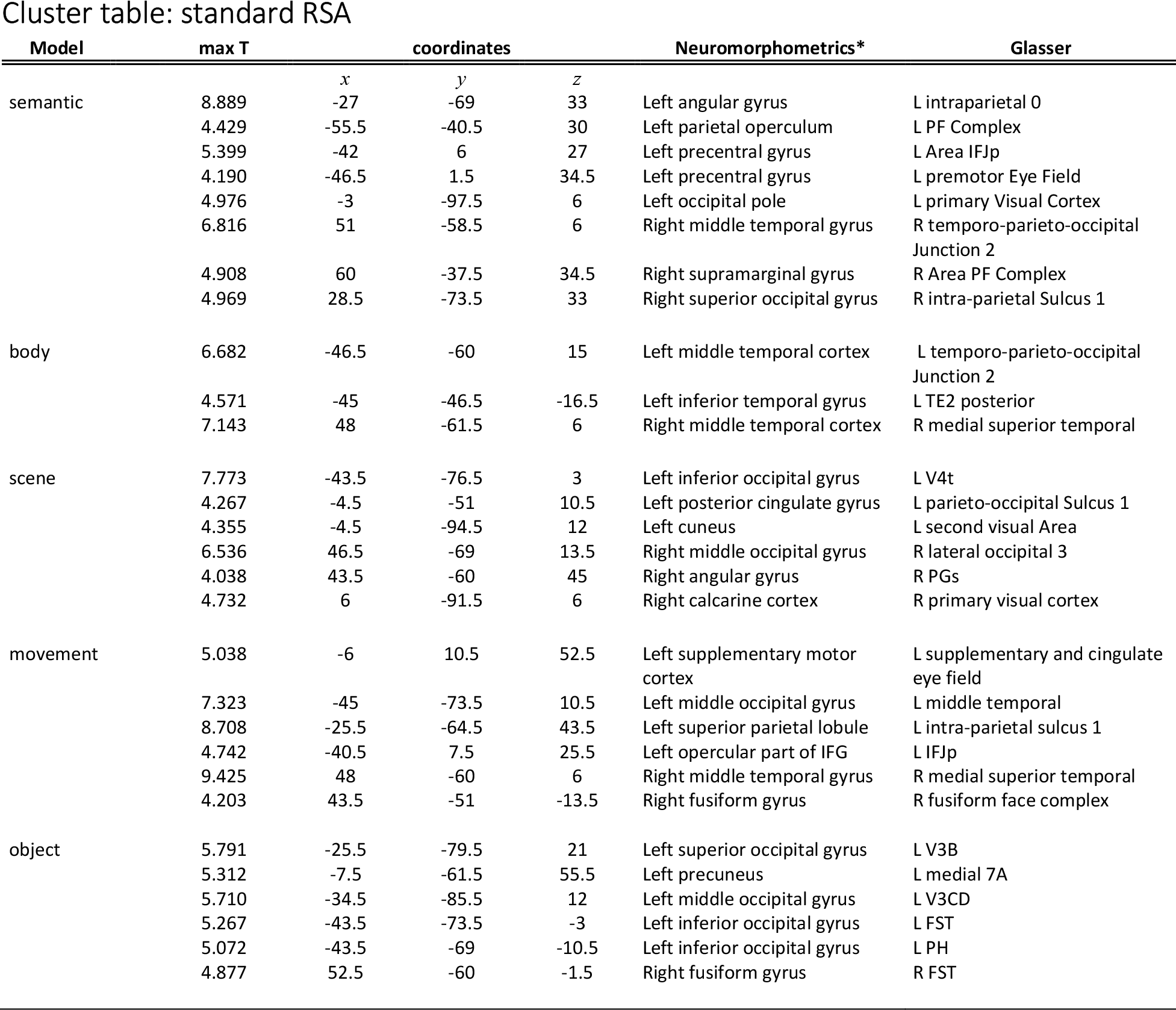
**List of clusters resulting from the standard RSA for the five different models** (semantic, body, scene, movement, object) which survived correction for multiple comparisons (cluster p-value < 0.05; see Methods and Figures 4 and 5). Coordinates are in MNI space. Labels are based on MRI scans that originated from the OASIS project (http://www.oasis-brains.org/) and were provided by Neuromorphometrics, Inc. (http://www.neuromorphometrics.com/) under academic subscription provided in SPM12 and the Glasser’s surface-based atlas^1^.

**Table S3.**
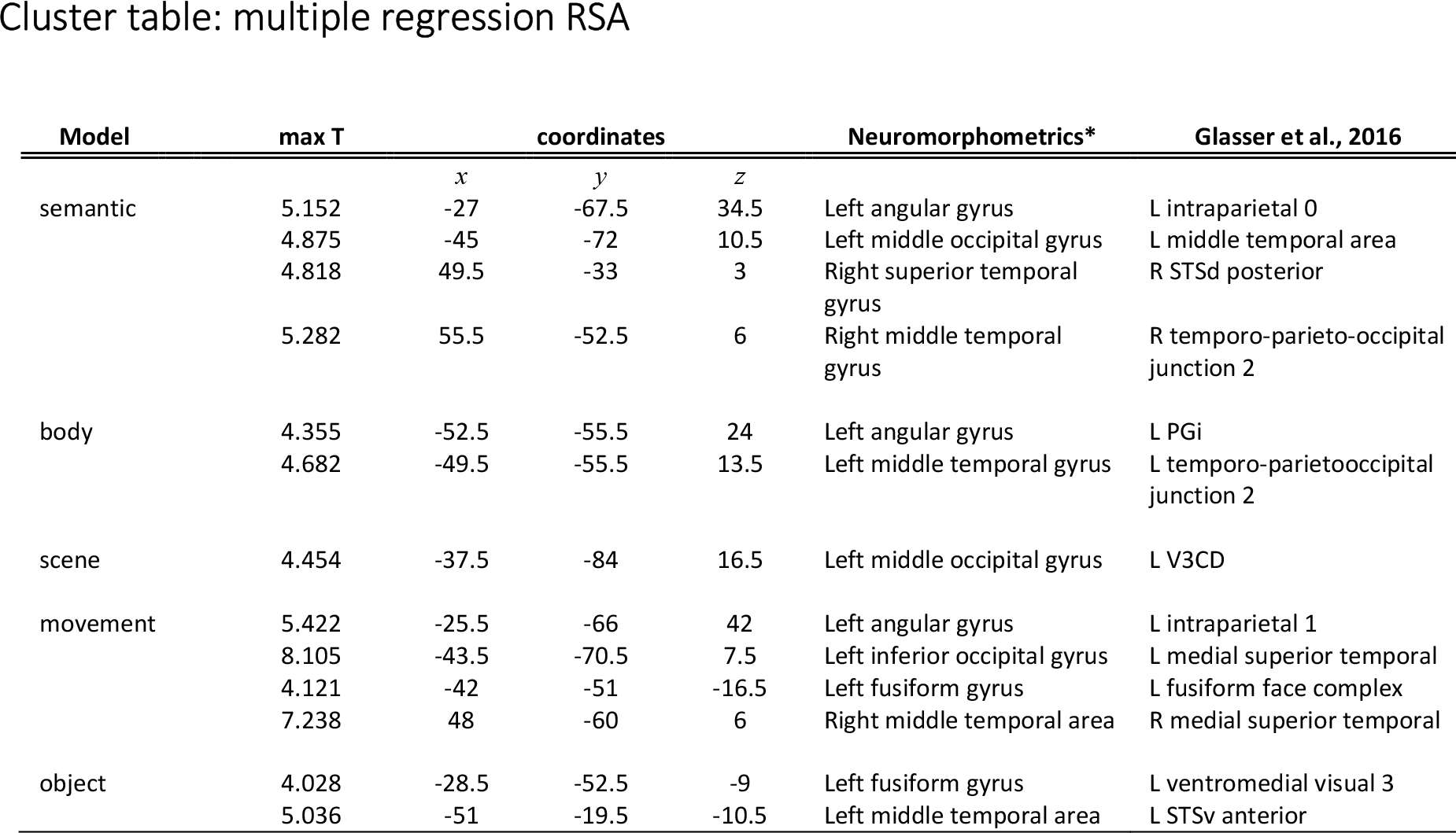
*List of clusters resulting from the multiple regression RSA for the five different models* (semantic, body, scene, movement, object) which survived correction for multiple comparisons (cluster p-value < 0.05; see Methods and Figures 6 and Figure S7). Coordinates are in MNI space. Labels are based on MRI scans that originated from the OASIS project (http://www.oasis-brains.org/) and were provided by Neuromorphometrics, Inc. (http://www.neuromorphometrics.com/) under academic subscription provided in SPM12 and the Glasser’s surface-based atlas^1^.

## References

1. Culham, J. C. & Valyear, K. F. Human parietal cortex in action. Curr. Opin. Neurobiol. 16, 205–212 (2006).

2. Rizzolatti, G. & Matelli, M. Two different streams form the dorsal visual system: anatomy and functions. Exp. brain Res. 153, 146–57 (2003).

3. Turella, L. & Lingnau, A. Neural correlates of grasping. Front. Hum. Neurosci. 8, 686 (2014).

4. Abdollahi, R. O., Jastorff, J. & Orban, G. A. Common and Segregated Processing of Observed Actions in Human SPL. Cereb. Cortex 23, 2734–2753 (2013).

5. Jastorff, J., Begliomini, C., Fabbri-Destro, M., Rizzolatti, G. & Orban, G. A. Coding Observed Motor Acts: Different Organizational Principles in the Parietal and Premotor Cortex of Humans. J. Neurophysiol. 104, 128–140 (2010).

6. Wurm, M. F., Caramazza, A. & Lingnau, A. Action Categories in Lateral Occipitotemporal Cortex Are Organized Along Sociality and Transitivity. J. Neurosci. 37, 562–575 (2017).

7. Haxby, J. V et al. Distributed and overlapping representations of faces and objects in ventral temporal cortex. Science 293, 2425–30 (2001).

8. Wurm, M. F., Ariani, G., Greenlee, M. W. & Lingnau, A. Decoding Concrete and Abstract Action Representations During Explicit and Implicit Conceptual Processing. Cereb. Cortex 26, 3390–3401 (2016).

9. Wurm, M. F. & Lingnau, A. Decoding Actions at Different Levels of Abstraction. J. Neurosci. 35, 7727–7735 (2015).

10. Hafri, A., Trueswell, J. C. & Epstein, R. A. Neural Representations of Observed Actions Generalize across Static and Dynamic Visual Input. J. Neurosci. 37, 3056–3071 (2017).

11. Oosterhof, N. N., Wiggett, A. J., Diedrichsen, J., Tipper, S. P. & Downing, P. E. Surface-based information mapping reveals crossmodal vision-action representations in human parietal and occipitotemporal cortex. J. Neurophysiol. 104, 1077–89 (2010).

12. Oosterhof, N. N., Tipper, S. P. & Downing, P. E. Viewpoint (in)dependence of action representations: an MVPA study. J. Cogn. Neurosci. 24, 975–89 (2012).

13. Bracci, S. & Peelen, M. V. Body and Object Effectors: The Organization of Object Representations in High-Level Visual Cortex Reflects Body-Object Interactions. J. Neurosci. 33, 18247–18258 (2013).

14. Kriegeskorte, N. & Mur, M. Inverse MDS: Inferring Dissimilarity Structure from Multiple Item Arrangements. Front. Psychol. 3, 1–13 (2012).

15. Kriegeskorte, N., Mur, M. & Bandettini, P. Representational similarity analysis – connecting the branches of systems neuroscience. Front. Syst. Neurosci. 2, 4 (2008).

16. Stelzer, J., Chen, Y. & Turner, R. Statistical inference and multiple testing correction in classification-based multi-voxel pattern analysis (MVPA): random permutations and cluster size control. Neuroimage 65, 69–82 (2013).

17. Maslow, a H. A theory of human motivation. Psychol. Rev. 50, 370–396 (1943).

18. Caspers, S., Zilles, K., Laird, A. R. & Eickhoff, S. B. ALE meta-analysis of action observation and imitation in the human brain. Neuroimage 50, 1148–1167 (2010).

19. Watson, C. E., Cardillo, E. R., Ianni, G. R. & Chatterjee, A. Action Concepts in the Brain: An Activation Likelihood Estimation Meta-analysis. J. Cogn. Neurosci. 25, 1191–1205 (2013).

20. Wurm, M. F. & Caramazza, A. Distinct roles of temporal and frontoparietal cortex in representing actions across vision and language. Nat. Commun. 10, 289 (2019).

21. Lingnau, A. & Downing, P. E. The lateral occipitotemporal cortex in action. Trends Cogn. Sci. (2015). doi:10.1016/j.tics.2015.03.006

22. Nasr, S. et al. Scene-selective cortical regions in human and nonhuman primates. J. Neurosci. 31, 13771–85 (2011).

23. Dilks, D. D., Julian, J. B., Paunov, A. M. & Kanwisher, N. The Occipital Place Area Is Causally and Selectively Involved in Scene Perception. J. Neurosci. 33, 1331–1336 (2013).

24. Kamps, F. S., Julian, J. B., Kubilius, J., Kanwisher, N. & Dilks, D. D. The occipital place area represents the local elements of scenes. Neuroimage (2016). doi:10.1016/j.neuroimage.2016.02.062

25. Bonner, M. F. & Epstein, R. A. Coding of navigational affordances in the human visual system. Proc. Natl. Acad. Sci. 114, 4793–4798 (2017).

26. Kim, H. J. et al. Canonical Correlation Analysis on Riemannian Manifolds and Its Applications. Eur. Conf. Comput. Vis. 251–267 (2014). doi:10.1007/978-3-319-10605-2_17

27. Greene, M. R., Baldassano, C., Esteva, A., Beck, D. M. & Fei-Fei, L. Visual scenes are categorized by function. J. Exp. Psychol. Gen. 145, 82–94 (2016).

28. El-Sourani, N., Wurm, M. F., Trempler, I., Fink, G. R. & Schubotz, R. I. Making sense of objects lying around: How contextual objects shape brain activity during action observation. Neuroimage 167, 429–437 (2018).

29. Bracci, S. & Op de Beeck, H. Dissociations and Associations between Shape and Category Representations in the Two Visual Pathways. J. Neurosci. 36, 432–444 (2016).

30. Kriegeskorte, N. et al. Matching categorical object representations in inferior temporal cortex of man and monkey. Neuron 60, 1126–41 (2008).

31. Chao, L. L., Haxby, J. V. & Martin, A. Attribute-based neural substrates in temporal cortex for perceiving and knowing about objects. Nat. Neurosci. (1999). doi:10.1038/13217

32. Caramazza, A. & Shelton, J. R. Domain-specifc knowledge systems in the brain: the animate-inanimate distinction. J. Cogn. Neurosci. 10, 1–34 (1998).

33. Mur, M. et al. Human Object-Similarity Judgments Reflect and Transcend the Primate-IT Object Representation. Front. Psychol. 4, 128 (2013).

34. Mecklinger, A., Gruenewald, C., Besson, M., Magnié, M.-N. & Von Cramon, D. Y. Separable neuronal circuitries for manipulable and non-manipulable objects in working memory. Cereb. Cortex 12, 1115–23 (2002).

35. New, J., Cosmides, L. & Tooby, J. Category-specific attention for animals reflects ancestral priorities, not expertise. Proc. Natl. Acad. Sci. U. S. A. 104, 16598–603 (2007).

36. Groen, I. I. A. et al. Distinct contributions of functional and deep neural network features to representational similarity of scenes in human brain and behavior. Elife 7, 1–26 (2018).

37. Schwarzbach, J. A simple framework (ASF) for behavioral and neuroimaging experiments based on the psychophysics toolbox for MATLAB. Behav. Res. Methods 43, 1194–201 (2011).

38. Brainard, D. H. The Psychophysics Toolbox. Spat. Vis. 10, 433–436 (1997).

39. Fischl, B., Sereno, M. I. & Dale, A. M. Cortical Surface-Based Analysis. Neuroimage 9, 195–207 (1999).

40. Oosterhof, N. N., Wiestler, T., Downing, P. E. & Diedrichsen, J. A comparison of volume-based and surface-based multi-voxel pattern analysis. Neuroimage 56, 593–600 (2011).

41. Oosterhof, N. N., Connolly, A. C. & Haxby, J. V. CoSMoMVPA: Multi-Modal Multivariate Pattern Analysis of Neuroimaging Data in Matlab/GNU Octave. Front. Neuroinform. 10, (2016).

42. Glasser, M. F. et al. The minimal preprocessing pipelines for the Human Connectome Project. Neuroimage 80, 105–124 (2013).

43. Marcus, D. S. et al. Informatics and Data Mining Tools and Strategies for the Human Connectome Project. Front. Neuroinform. 5, 1–12 (2011).

44. Hartigan, J. A. & Wong, M. A. A K-Means Clustering Algorithm. Appl. Stat. 28, 100 (1979).

45. Rousseeuw, P. J. Silhouettes: A graphical aid to the interpretation and validation of cluster analysis. J. Comput. Appl. Math. 20, 53–65 (1987).

46. Kriegeskorte, N., Goebel, R. & Bandettini, P. Information-based functional brain mapping. Proc. Natl. Acad. Sci. 103, 3863–3868 (2006).

47. Mason, R. L., Gunst, R. F. & Hess, J. L. Statistical Design and Analysis of Experiments. (John Wiley & Sons, Inc., 2003). doi:10.1002/0471458503

## SI References

1. Glasser, MF, et al. (2016) A multi-modal parcellation of human cerebral cortex. Nature 536(7615):171–178.

