## supplementary information for "The representational space of observed actions"

#### Images of the actions used in the fMRI experiment

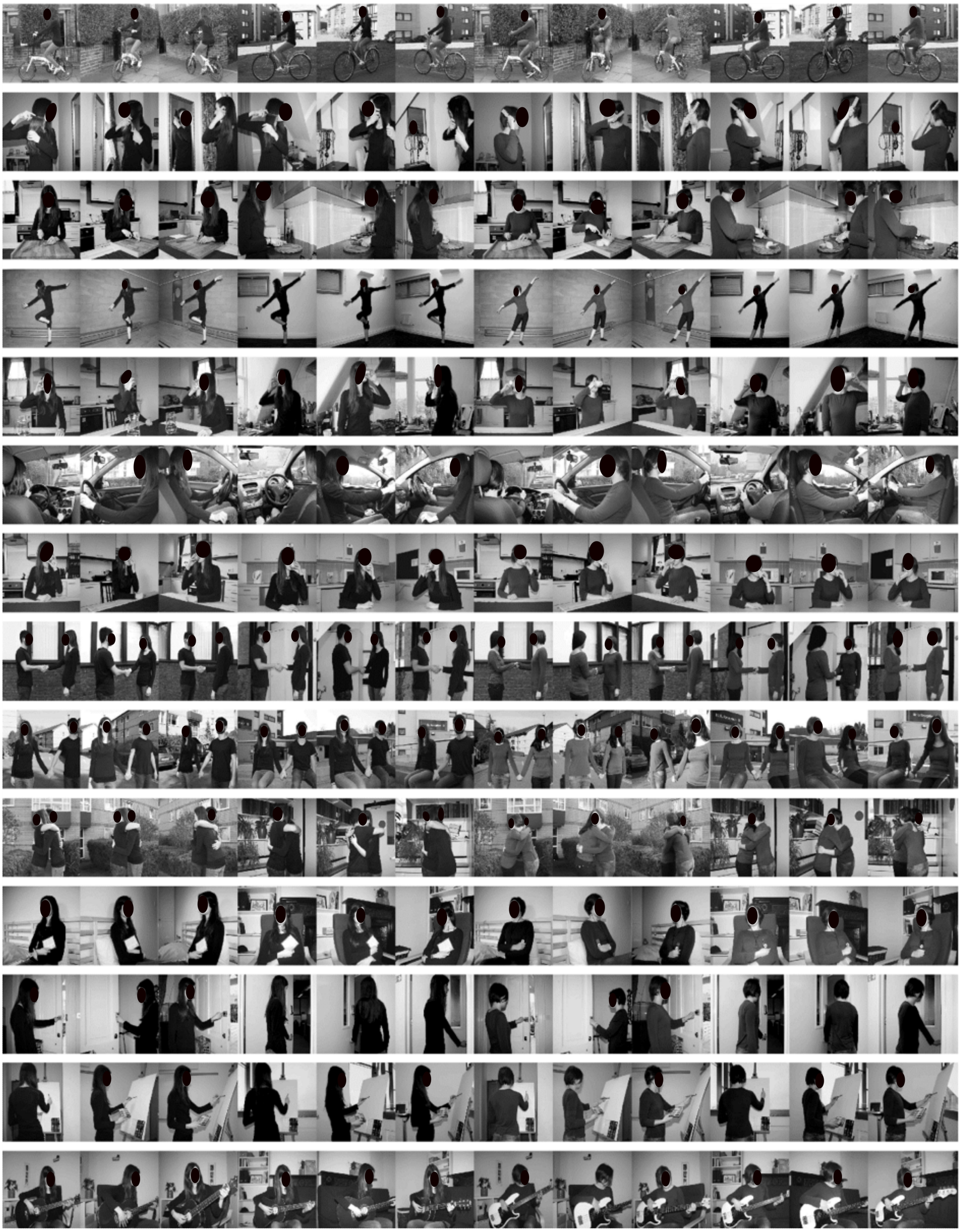

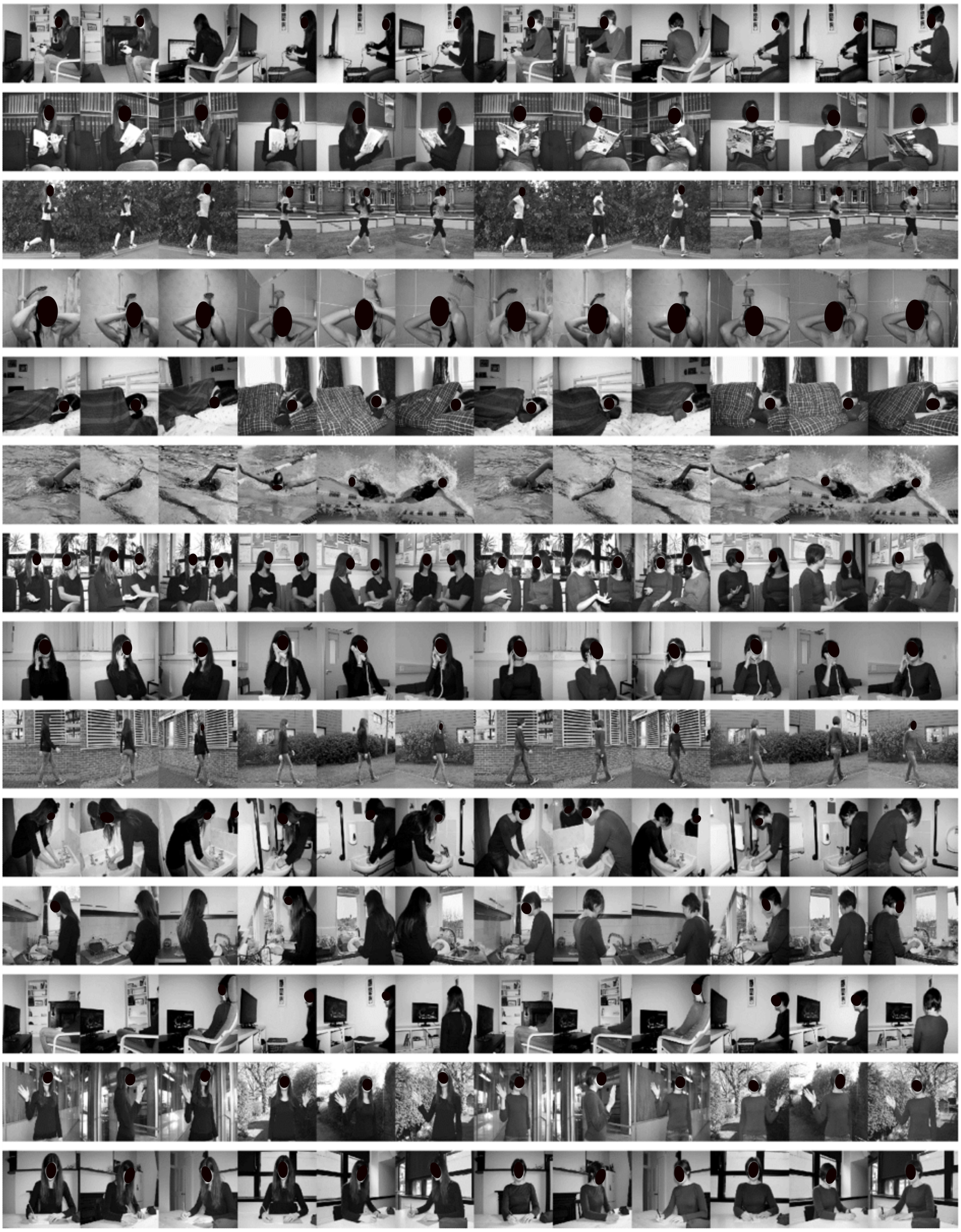

***Figure S1.*** *All images used in the fMRI experiment. Different actions are depicted in rows (for corresponding labels, see words printed in bold font in* ***Table S1****). Different exemplars for each action are shown in the columns. Exemplars varied in terms of the actor (2), scene (2), and viewing angle (3).*

#### Pairwise correlation between models averaged across participants

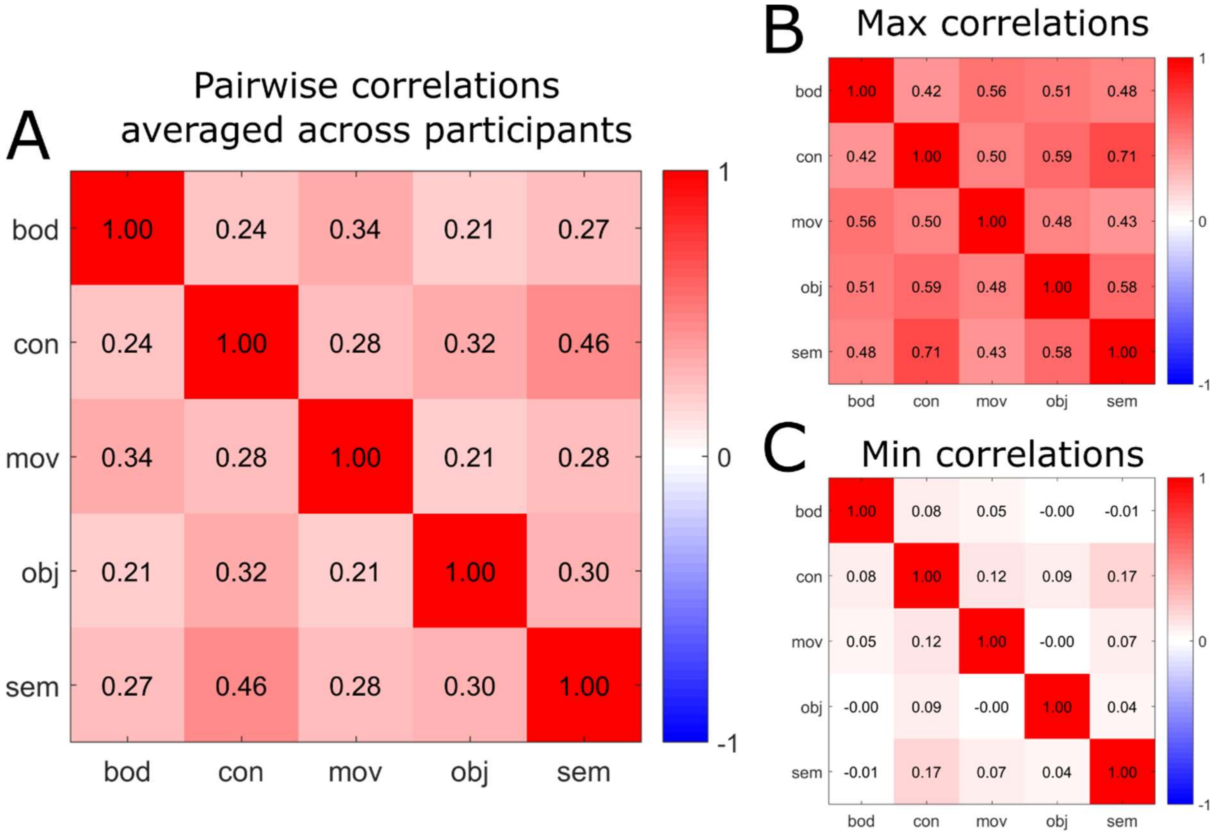


***Figure S2.*** *Pairwise cross-correlation matrix across models. For each participant, we computed the pairwise cross-correlation matrix across models. We then computed (A) the averaged correlation values across participants; and reporting (B) the maximum and (C) the minimum correlation values across participants. For a corresponding analysis of collinearity using the variance inflation factor, see Methods, Multiple Regression RSA.*

#### Silhouette analysis

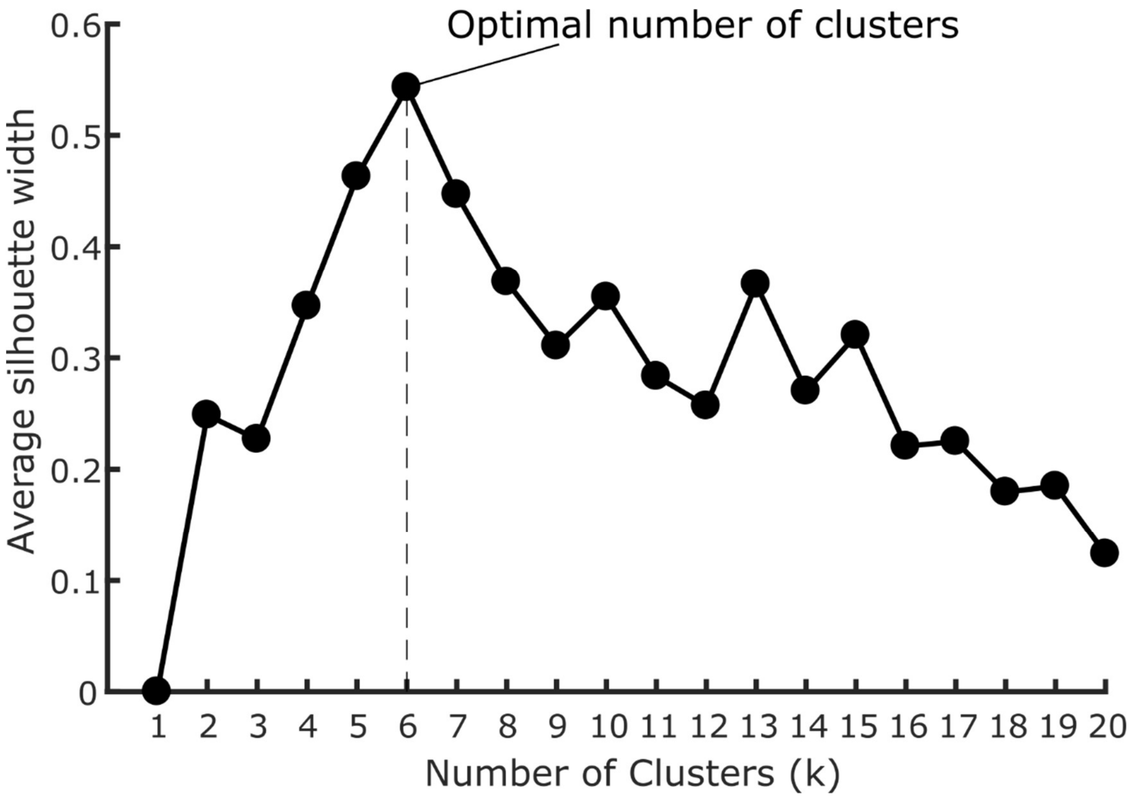


***Figure S3.*** *Silhouette analysis revealed that the optimal number of clusters for the Semantic model was 6. The analysis was performed using the fviz_nbclust function of the R package "factoextra". The silhouette width is a measure of how well each item fits within its cluster with higher values indicating better fit. The average silhouette width is the average of the all items. The algorithm computes the average silhouette width as a function of the number of clusters (k). The highest value of the average silhouette width is taken and the corresponding k is selected as the optimal number of clusters.*

#### Eigenvalues of the Principal Component Analysis

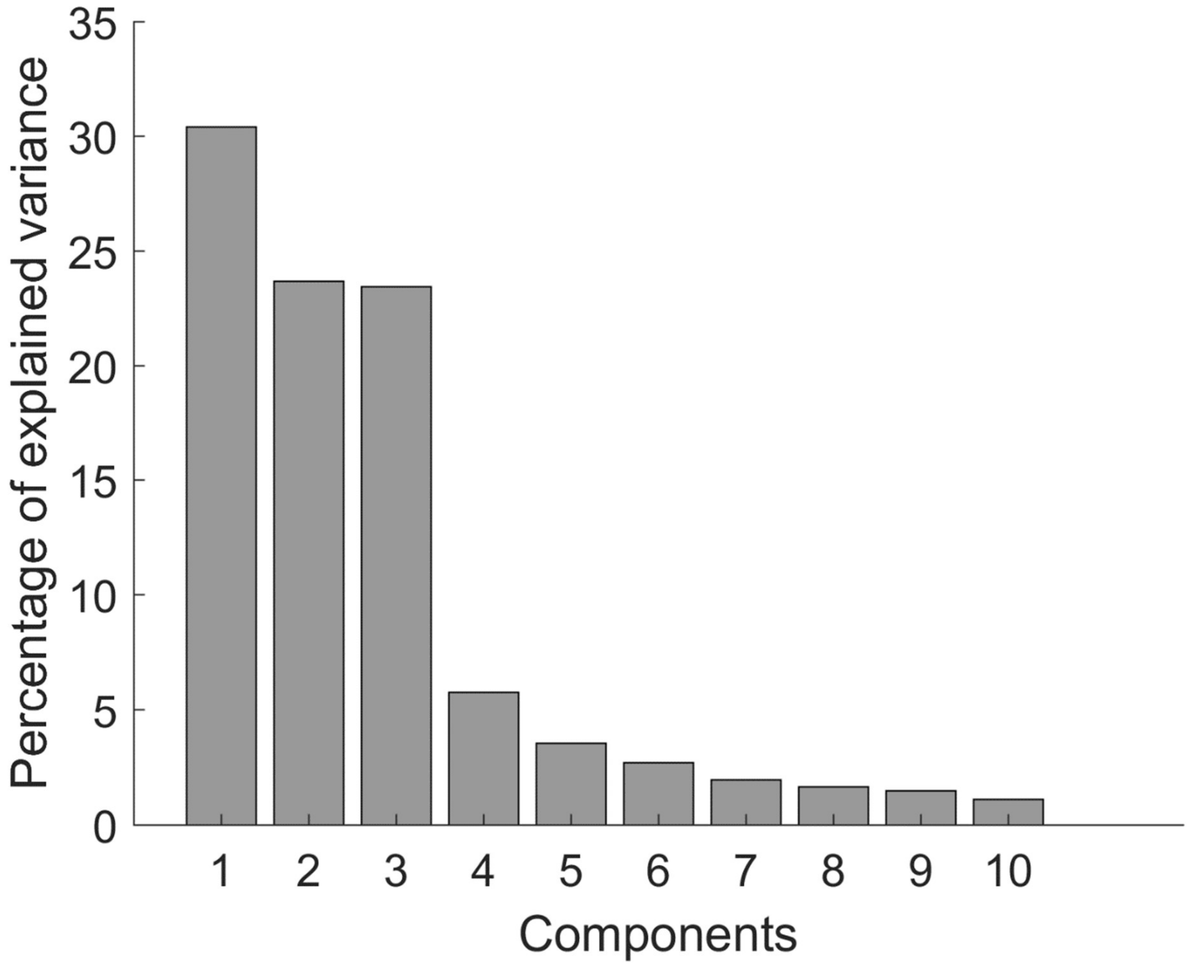


***Figure S4.*** *Eigenvalues obtained from the PCA of the semantic model. The PCA shows that the first 3 components accounted for the largest amount of variance*

#### PCA and K-means

semantic model

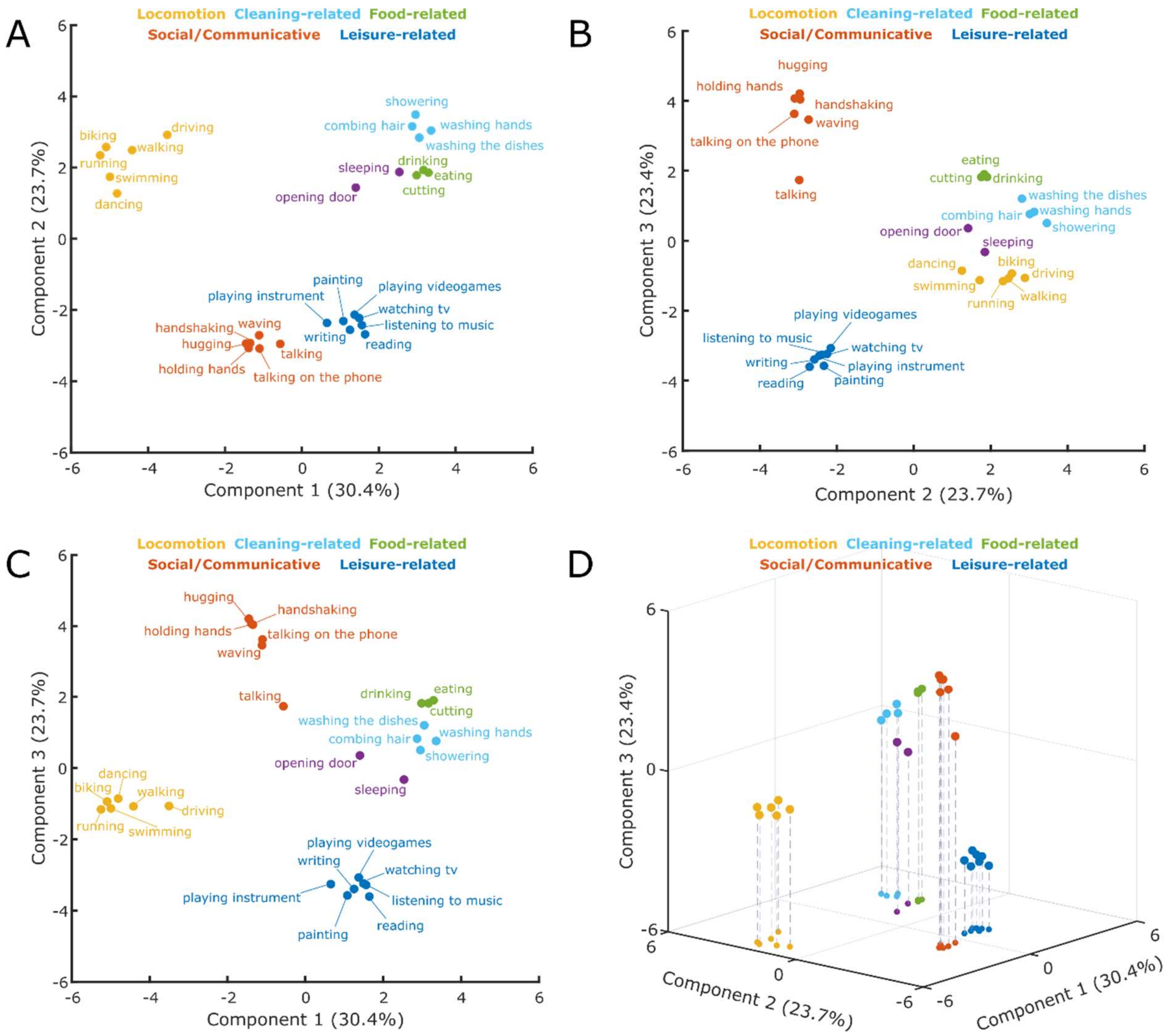


***Figure S5.*** *Cluster analysis. Clusters resulting from the K-means clustering analysis (K-means) for the semantic model. A: 2D-plot showing component 1 and 2, corresponding labels of individual actions and suggested labels for the categories resulting from the K-means clustering. B: 2D-plot showing component 2 and 3. C: 2D-plot showing component 1 and 3. D: 3D-plot showing components 1-3.*

#### Other models

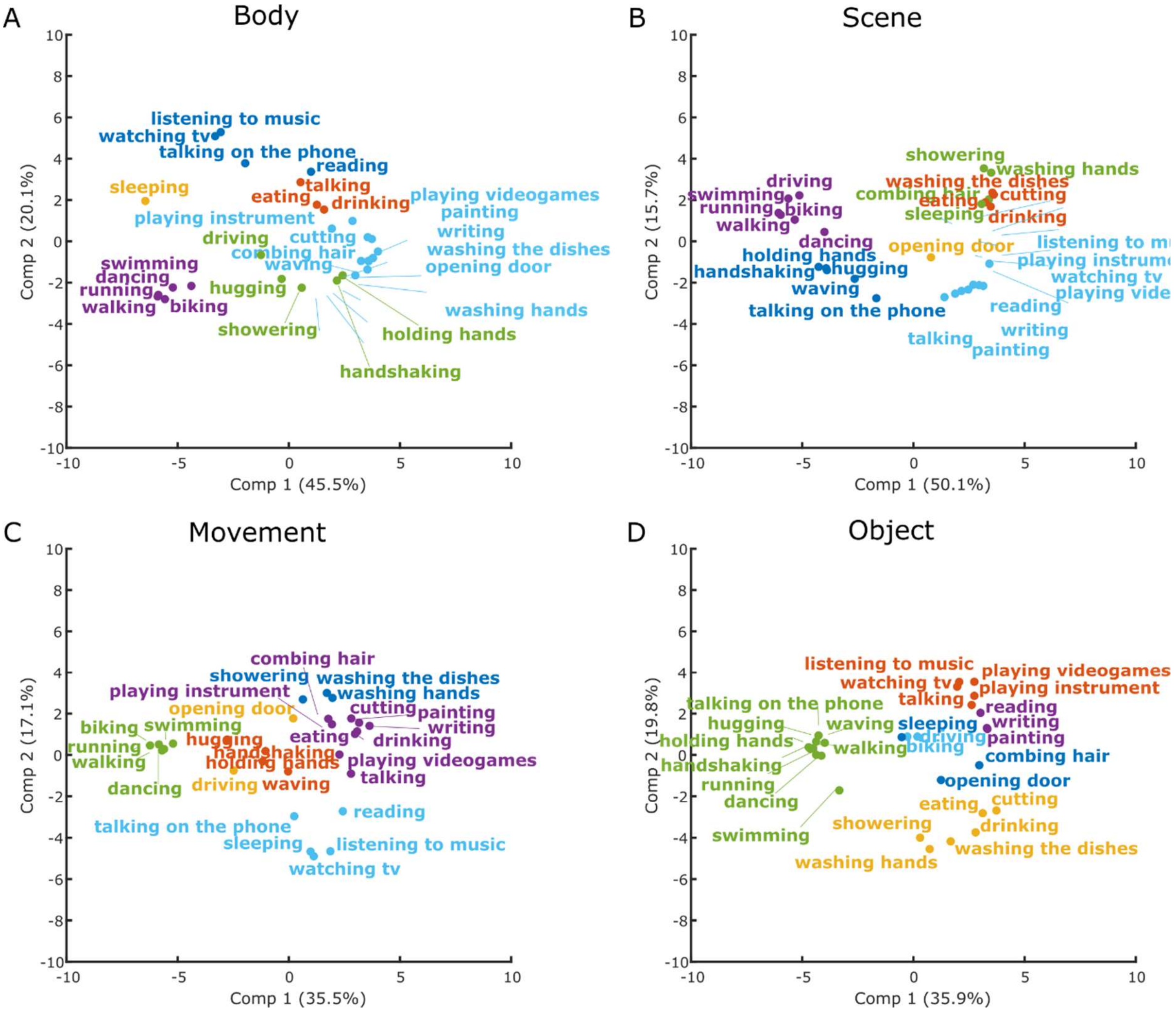


***Figure S6.*** *First 2 principal components of the control models. Clusters were distinguished using the K-Means methods as implemented in R. Number of clusters used as input were 6 (based on the Silhouette analyses of the semantic model; see Figure S3). Colors are used to distinguish between the six resulting clusters in each analysis.*

### Multiple regression RSA for the control models

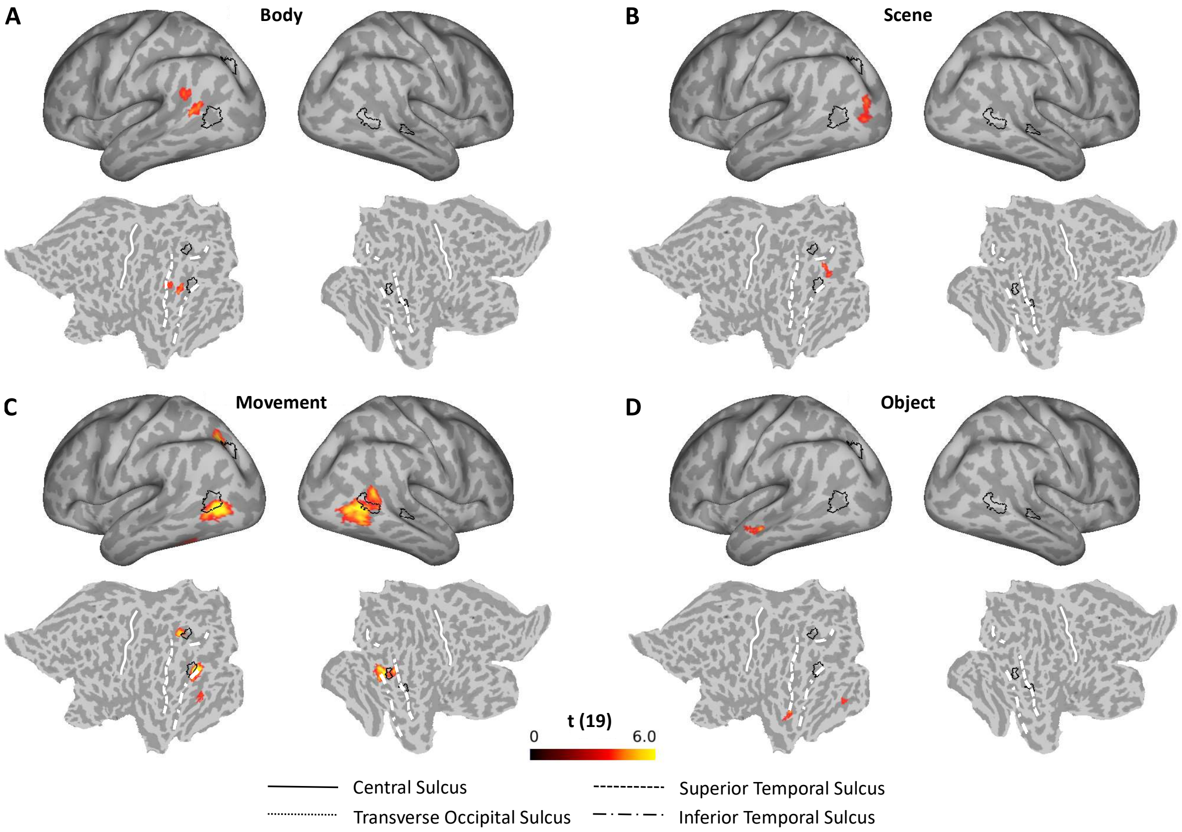


***Figure S7.*** *Multiple regression RSA, comparison between the semantic and the other models. Group results of the searchlight-based multiple regression RSA for the body (A), scene (B), movement (C), and object (D) model. For details, see methods section and captions of Figure 4. Black outlines on the inflated brains and the flat maps depict significant clusters revealed by the multiple regression RSA for the semantic mode (Figure 6).*

#### List of actions

| Actions | | | |
|---|---|---|---|
| **biking** | grocery shopping | **reading** | **walking** |
| brushing teeth | **handshaking** | **running** | **washing hands** |
| **combing hair** | **holding hands** | singing | **washing the dishes** |
| cleaning the floor | **hugging** | **sleeping** | **watching tv** |
| **cutting** | **listening to music** | **swimming** | watering plants |
| **dancing** | making coffee | switching on light | **waving** |
| **drinking** | **opening door** | **taking a shower** | **writing** |
| **driving** | **painting** | taking the train | |
| **eating** | **playing instrument** | **talking on the phone** | |
| getting dressed | **playing videogames** | **talking** | |

***Table S1.*** *List of actions identified using the online survey. We initially kept actions that were mentioned by at least 20% of the participants. The final selected twenty-eight actions used for the study are the ones highlighted. See the Methods section of the main article for the procedure we used to select these actions.*

#### Cluster table: standard RSA

| Model | max T | coordinates | | | Neuromorphometrics* | Glasser |
|---|---|---|---|---|---|---|
| | | *x* | *y* | *z* | | |
| semantic | 8.889 | -27 | -69 | 33 | Left angular gyrus | L intraparietal 0 |
| | 4.429 | -55.5 | -40.5 | 30 | Left parietal operculum | L PF Complex |
| | 5.399 | -42 | 6 | 27 | Left precentral gyrus | L Area IFJp |
| | 4.190 | -46.5 | 1.5 | 34.5 | Left precentral gyrus | L premotor Eye Field |
| | 4.976 | -3 | -97.5 | 6 | Left occipital pole | L primary Visual Cortex |
| | 6.816 | 51 | -58.5 | 6 | Right middle temporal gyrus | R temporo-parieto-occipital Junction 2 |
| | 4.908 | 60 | -37.5 | 34.5 | Right supramarginal gyrus | R Area PF Complex |
| | 4.969 | 28.5 | -73.5 | 33 | Right superior occipital gyrus | R intra-parietal Sulcus 1 |
| body | 6.682 | -46.5 | -60 | 15 | Left middle temporal cortex | L temporo-parieto-occipital Junction 2 |
| | 4.571 | -45 | -46.5 | -16.5 | Left inferior temporal gyrus | L TE2 posterior |
| | 7.143 | 48 | -61.5 | 6 | Right middle temporal cortex | R medial superior temporal |
| scene | 7.773 | -43.5 | -76.5 | 3 | Left inferior occipital gyrus | L V4t |
| | 4.267 | -4.5 | -51 | 10.5 | Left posterior cingulate gyrus | L parieto-occipital Sulcus 1 |
| | 4.355 | -4.5 | -94.5 | 12 | Left cuneus | L second visual Area |
| | 6.536 | 46.5 | -69 | 13.5 | Right middle occipital gyrus | R lateral occipital 3 |
| | 4.038 | 43.5 | -60 | 45 | Right angular gyrus | R PGs |
| | 4.732 | 6 | -91.5 | 6 | Right calcarine cortex | R primary visual cortex |
| movement | 5.038 | -6 | 10.5 | 52.5 | Left supplementary motor cortex | L supplementary and cingulate eye field |
| | 7.323 | -45 | -73.5 | 10.5 | Left middle occipital gyrus | L middle temporal |
| | 8.708 | -25.5 | -64.5 | 43.5 | Left superior parietal lobule | L intra-parietal sulcus 1 |
| | 4.742 | -40.5 | 7.5 | 25.5 | Left opercular part of IFG | L IFJp |
| | 9.425 | 48 | -60 | 6 | Right middle temporal gyrus | R medial superior temporal |
| | 4.203 | 43.5 | -51 | -13.5 | Right fusiform gyrus | R fusiform face complex |
| object | 5.791 | -25.5 | -79.5 | 21 | Left superior occipital gyrus | L V3B |
| | 5.312 | -7.5 | -61.5 | 55.5 | Left precuneus | L medial 7A |
| | 5.710 | -34.5 | -85.5 | 12 | Left middle occipital gyrus | L V3CD |
| | 5.267 | -43.5 | -73.5 | -3 | Left inferior occipital gyrus | L FST |
| | 5.072 | -43.5 | -69 | -10.5 | Left inferior occipital gyrus | L PH |
| | 4.877 | 52.5 | -60 | -1.5 | Right fusiform gyrus | R FST |

***Table S2. List of clusters resulting from the standard RSA for the five different models*** *(semantic, body, scene, movement, object) which survived correction for multiple comparisons (cluster p-value < 0.05; see Methods and Figures 4 and 5). Coordinates are in MNI space. Labels are based on MRI scans that originated from the OASIS project (http://www.oasis-brains.org/) and were provided by Neuromorphometrics, Inc. (http://www.neuromorphometrics.com/) under academic subscription provided in SPM12 and the Glasser's surface-based atlas[1].*

#### Cluster table: multiple regression RSA

| Model | max T | coordinates | | | Neuromorphometrics* | Glasser et al., 2016 |
|---|---|---|---|---|---|---|
| | | *x* | *y* | *z* | | |
| semantic | 5.152 | -27 | -67.5 | 34.5 | Left angular gyrus | L intraparietal 0 |
| | 4.875 | -45 | -72 | 10.5 | Left middle occipital gyrus | L middle temporal area |
| | 4.818 | 49.5 | -33 | 3 | Right superior temporal gyrus | R STSd posterior |
| | 5.282 | 55.5 | -52.5 | 6 | Right middle temporal gyrus | R temporo-parieto-occipital junction 2 |
| body | 4.355 | -52.5 | -55.5 | 24 | Left angular gyrus | L PGi |
| | 4.682 | -49.5 | -55.5 | 13.5 | Left middle temporal gyrus | L temporo-parietooccipital junction 2 |
| scene | 4.454 | -37.5 | -84 | 16.5 | Left middle occipital gyrus | L V3CD |
| movement | 5.422 | -25.5 | -66 | 42 | Left angular gyrus | L intraparietal 1 |
| | 8.105 | -43.5 | -70.5 | 7.5 | Left inferior occipital gyrus | L medial superior temporal |
| | 4.121 | -42 | -51 | -16.5 | Left fusiform gyrus | L fusiform face complex |
| | 7.238 | 48 | -60 | 6 | Right middle temporal area | R medial superior temporal |
| object | 4.028 | -28.5 | -52.5 | -9 | Left fusiform gyrus | L ventromedial visual 3 |
| | 5.036 | -51 | -19.5 | -10.5 | Left middle temporal area | L STSv anterior |

***Table S3. List of clusters resulting from the multiple regression RSA for the five different models*** *(semantic, body, scene, movement, object) which survived correction for multiple comparisons (cluster p-value < 0.05; see Methods and Figures 6 and Figure S7). Coordinates are in MNI space. Labels are based on MRI scans that originated from the OASIS project (http://www.oasis-brains.org/) and were provided by Neuromorphometrics, Inc. (http://www.neuromorphometrics.com/) under academic subscription provided in SPM12 and the Glasser's surface-based atlas[1].*

#### SI References

1. Glasser MF, et al. (2016) A multi-modal parcellation of human cerebral cortex. *Nature* 536(7615):171–178.
